# The DUF998 family protein DrmA modulates cationic glycolipid levels and the emergence of high-level daptomycin resistance in *Enterococcus faecalis*

**DOI:** 10.64898/2026.07.07.736841

**Authors:** Aparna Uppuluri, Jonathan Martin, Luke R. Joyce, Tata Ninidze, Kelly S. Doran, Faruck Morcos, Ziqiang Guan, Kelli L. Palmer

## Abstract

Daptomycin resistance (DAP-R) in enterococci is associated with alterations in the membrane lipid composition. The membrane-bound protein MprF is responsible for the synthesis of amino acid-modified lipids in bacteria, and these modified lipids contribute to DAP-R in some Gram-positive pathogens. In enterococci, MprF synthesizes three lysine-modified lipids: the phospholipid lysyl-phosphatidylglycerol (Lys-PG), and the newly identified cationic glycolipids lysyl-diglucosyl-diacylglycerol (Lys-Glc_2_-DAG) and lysyl-glucosyl-diacylglycerol (Lys-Glc-DAG). Given the recent discovery of cationic glycolipids in enterococci, we re-examined a collection of laboratory-evolved DAP-R *E. faecalis* to investigate whether these lipids contribute to DAP-R. We found that levels of Lys-Glc_2_-DAG were strikingly reduced in DAP-R variants with high-level resistance. The dramatic alterations in Lys-Glc_2_-DAG levels were temporally coupled with the emergence of loss-of-function mutations in the gene *drmA*, which encodes a DUF998 family protein of unknown function. DrmA is a membrane protein with six predicted transmembrane helices and is widely distributed among Gram-positive and Gram-negative bacteria, including plant and animal pathogens. Complementation of the DAP-R strains with wild-type *E. faecalis drmA* significantly lowered their DAP MIC, reversing their trajectory to high-level DAP-R. Using genetic and lipidomic approaches in the natively DAP-sensitive strain OG1RF, we conclusively linked *drmA* loss-of-function with significantly reduced Lys-Glc_2_-DAG levels as well as a small but significant increase in Lys-PG levels. Yet, *drmA* inactivation in OG1RF did not alter its DAP MIC. We conclude that *drmA* loss-of-function confers elevated DAP MIC on the background of preceding mutations in the DAP-R evolutionary trajectory, most likely mutations in *cls1*. The recurrence of *drmA* mutations in multiple studies underscores its importance in DAP-R evolution. Overall, our work identifies a role for the DUF998 family in cellular lipid homeostasis and confirms its significant role in the evolution of DAP-R.

## Introduction

*Enterococcus faecalis* and *Enterococcus faecium* are Gram-positive bacteria and human gut commensals that can cause devastating infections, including bacteremia and infective endocarditis (1). Enterococci have robust and adaptable cell surfaces (2). The enterococcal cell surface is comprised of peptidoglycan, cell surface polymers (including teichoic acids and capsule), lipids, and proteins,-all of which play important roles in maintaining cell function and structural integrity (3, 4).

Among these cell surface components, membrane lipids in particular have important roles in enterococcal host-pathogen interactions and antimicrobial resistance (5, 6). The major lipids of the enterococcal membrane include the phospholipids phosphatidylglycerol (PG) and its derivatives cardiolipin (CL) and lysyl-phosphatidylglycerol (Lys-PG), and the glycolipids monoglucosyl-diacylglycerol (Glc-DAG) and diglucosyl-diacylglycerol (Glc_2_-DAG) (5, 7, 8). In addition to their roles in forming the membrane, PG and glucosyl-DAGs also serve as substrates for the synthesis of the membrane-anchored polymer Type I lipoteichoic acid (LTA) and for cationic phospholipids and glycolipids (aminoacylated lipids, discussed below) (5, 7–11).

The protein MprF (Multiple Peptide Resistance Factor) is extensively studied for its role in modifying lipid head groups with amino acids, thereby allowing bacteria to alter their surface charge. More specifically, the aminoacylation of PG by MprF neutralizes the negative charge of PG, aids in bacterial virulence, and confers protection against cationic antimicrobial peptides (CAMPs) (12, 13). Interestingly, MprF proteins from different organisms may have different catalytic activities, including the use of lipid substrates other than PG and the incorporation of different amino acids for lipid modification, most commonly lysine and alanine (7, 9). *E. faecalis* and *E. faecium* encode two *mprF* paralogs, *mprF1* and *mprF2* (7); note that *mprF1* in *E. faecalis* is the ortholog of *mprF2* in *E. faecium*, whereas *mprF2* in *E. faecalis* is the ortholog of *mprF1* in *E. faecium*. Recently, we discovered that *E. faecalis* MprF2 catalyzes the synthesis of three distinct lysine-modified lipids, namely the novel cationic glycolipids lysyl-diglucosyl-diacylglycerol (Lys-Glc_2_-DAG) and lysyl-glucosyl-diacylglycerol (Lys-Glc-DAG), as well as Lys-PG, the well-known lipid observed in many other Gram-positive pathogens(9). The novel cationic glycolipids may play key roles in enterococcal antimicrobial resistance and host-pathogen interactions, but this has not been investigated due to their recent discovery.

Cationic lipids are associated with resistance to the cyclic lipopeptide antibiotic daptomycin (DAP), which is used as a last resort to treat Gram-positive bloodstream and skin and soft tissue infections (14, 15). One model for the DAP mechanism of action involves calcium ions and DAP forming a complex with PG and undecaprenyl phosphate (C_55_-P) cell wall precursors, leading to membrane disruption and cell lysis (16). Alarmingly, enterococci and other Gram-positive pathogens can develop resistance to DAP by mutation, leading to clinical treatment failures (17). In *S. aureus*, DAP-resistant (DAP-R) strains exhibit altered lipidomes characterized by increased Lys-PG levels, and the acquisition of DAP-R is often associated with mutations in *mprF* (18). In enterococci, DAP resistance has been linked to mutations in multiple genes, including the *liaFSR* regulatory system, the cardiolipin synthase gene *cls1*, and the presumptive loss-of-function mutations in a gene of unknown function referred to as *drmA* (gene locus EF1797 in the model *E. faecalis* strain V583) (8, 19–23).

Given our recent discovery of novel, highly cationic glycolipids in enterococci (9), which were not considered in prior models for DAP resistance emergence, we revisited a prior study of DAP resistance in *E. faecalis*. Three highly DAP-resistant (DAP-R) derivatives of the bloodstream infection isolate *E. faecalis* V583 were previously generated by *in vitro* evolution (19). Genome sequencing revealed mutations in seven candidate genes across these strains, including *drmA* (19). Here, we show that these high-level DAP-R strains possess unique lipid profiles compared to the parental V583 strain, specifically having strikingly lower levels of Lys-Glc_2_-DAG. We determined that DrmA regulates Lys-Glc_2_-DAG levels and that loss of *drmA* function confers high-level DAP resistance in *E. faecalis* V583, which can be reversed by restoration of *drmA*. DrmA belongs to the domain of unknown function (DUF) family DUF998, which is widely distributed in prokaryotes and is structurally analogous to the eukaryotic family of transmembrane protein family FRAG1/DRAM/sfk1 with proposed roles in establishing trans-bilayer asymmetry of phospholipids (24, 25). Overall, our work suggests a key role for cationic glycolipids in enterococcal-DAP interactions and establishes DrmA as a strong modulator of cationic glycolipid levels in enterococci.

## Materials and Methods

### Bacterial strains and routine growth conditions

The bacterial strains used in this study are listed in Table 1, Table 2, and Dataset S1. All strains in Table 1 are from previously published work (19). *E. faecalis* OG1RF transposon (Tn) mutants were obtained from a previously reported Tn mutant library (26). All *E. faecalis* OG1RF Tn mutants used in this study were verified by PCR with primers listed in Dataset S1. All enterococcal strains were grown at 37°C on brain heart infusion (BHI) agar or in BHI broth. Tn mutants and strains containing plasmids were grown in the presence of appropriate antibiotics (15 μg/mL chloramphenicol for Tn mutants (26) and either 1000 μg/mL kanamycin or 15 μg/ml chloramphenicol for the strains with pABG5Δ*phoZ* (27)). All enterococcal strains were cultured without agitation unless otherwise stated. *Escherichia coli* DH5*α* strains with pABG5Δ*phoZ* and its derivatives were grown on Luria-Bertani (LB) agar plates or in LB broth supplemented with kanamycin (100 μg/mL) at 37°C with agitation at 225 rpm. *Bacillus subtilis* ATCC 23857 was grown on LB agar plates or in LB broth at 37°C with agitation at 225 rpm.

**Table 1:** *E. faecalis* V583 DAP-R strains obtained by serial passage in a prior study.

| *E. faecalis* strain<sup>a</sup> | DAP MIC (μg/mL)<sup>b</sup> | *clsI* EF0631<sup>c</sup> | *drmA* EF1797<sup>c</sup> | EF0716<sup>c</sup> | *rpoN* EF0782<sup>c</sup> | *liaX* EF1753<sup>c</sup> | EF2470<sup>c</sup> | *telA* EF2698<sup>c</sup> |
| --- | --- | --- | --- | --- | --- | --- | --- | --- |
| V583 (Parent) | 4 | - | - | - | - | - | - | - |
| DAP-A Day 4 | 24-32 | N77_Q79 del | D30fs | - | - | - | - | - |
| DAP-A Day 12 | 96-128 | N77_Q79 del | D30fs | W148_F150dup | T99R | - | - | - |
| DAP-B Day 13 | 96-128 | R218Q | V16ins IS256 | - | - | 1390fs | I117fs | I277F |
| DAP-C Day 5 | 12-24 | R218Q | - | - | - | 1390fs | - | - |
| DAP-C Day 14 | >256 | R218Q | G130_F154del | W148_F150del | - | 1390fs | W82X | - |

<sup>a</sup>Strains and amino acid substitution data were previously published by Palmer, et al. (2011). Briefly, *E. faecalis* V583 was serially passaged in increasing concentrations of DAP for up to 14 days. Isolates from the serial passage are shown in this table, along with their day of isolation, DAP MIC, and mutations relative to the parental strain.
<sup>b</sup>DAP MIC was determined by E-test for all strains (n=3 independent replicates).
<sup>c</sup>Amino acid substitutions are shown for V583 gene loci. Abbreviations: del, deletion; fs, frameshift; dup, duplication; ins, insertion. Predicted functions of the affected V583 gene loci are as follows: EF1797 (*drmA*), DUF998 domain containing protein; EF0716 (*clsI*), cardiolipin synthetase; EF0716, hypothetical protein; EF0782 (*rpoN*), DNA-directed RNA polymerase sigma subunit RpoN; EF1753 (*liaX*), daptomycin-sensing surface protein LiaX; EF2470, HD domain-containing protein; and EF2698 (*telA*), tellurite resistance protein.

### Daptomycin E-test

DAP minimum inhibitory concentration (MIC) was assessed using Liofilchem DAP E-test strips according to the manufacturer’s instructions. The DAP E-test was performed in three independent replicates for all strains.

### Lipid Extraction

Lipid extractions of bacterial cultures were performed by acidic Bligh-Dyer lipid extraction as previously described (9, 28, 29). Unless otherwise noted, 14-15 mL of overnight culture was pelleted by centrifugation, washed twice with 1X phosphate buffered saline (PBS), and transferred to glass tubes for lipid extraction. 1 mL of chloroform and 2 mL methanol were added to each tube to make a single-phase Bligh-Dyer mixture, followed by centrifugation at 500×g for 10 min. The supernatant was transferred to a new glass tube and 100 μl of hydrochloric acid, 1 mL of chloroform and 0.9 mL of 1X PBS were added to make a two-phase Bligh-Dyer and centrifuged at 500×g for 5 minutes. The lower organic phase was transferred to a new glass tube and dried under a stream of nitrogen gas.

### Lipidomic analysis by liquid chromatography/electrospray ionization mass spectrometry

An amino column-based normal phase LC-ESI/MS was performed using an Agilent 1200 Quaternary LC system coupled to a high-resolution TripleTOF5600 mass spectrometer (Sciex, Framingham, MA). A Unison UK-Amino column (3 *μ*m, 25 cm × 2 mm) (Imtakt USA, Portland, OR) was used. Mobile phase A consisted of chloroform/methanol/aqueous ammonium hydroxide (800:195:5, v/v/v). Mobile phase B consisted of chloroform/methanol/water/ aqueous ammonium hydroxide (600:340:50:5, v/v/v/v). Mobile phase C consisted of chloroform/methanol/water/aqueous ammonium hydroxide (450:450:95:5, v/v/v/v). The elution program consisted of the following: 100% mobile phase A was held isocratically for 2 min and then linearly increased to 100% mobile phase B over 8 min and held at 100% B for 5 min. The LC gradient was then changed to 100% mobile phase C over 1 min and held at 100% C for 3 min, and finally returned to 100% A over 0.5 min and held at 100% A for 3 min. The total LC flow rate was 300 *μ*l/min. The MS settings were as follows: Ion spray voltage (IS) = +5000 V, Curtain gas (CUR) = 20 psi, Ion source gas 1 (GS1) = 20 psi, Declustering potential (DP) = +50 V, and Focusing Potential (FP) = +150 V. Nitrogen was used as the collision gas for MS/MS experiments. Data acquisition and analysis were performed using Analyst TF1.5 software (Sciex, Framingham, MA).

### Thin layer chromatography (TLC)

Cell pellets from 100 mL of overnight (stationary phase) cultures of *E. faecalis* were used for lipid extractions, with three independent replicates for each strain. The acidic Bligh-Dyer lipid extractions were performed as described above. 100 μL chloroform was added in order to resuspend dried, extracted lipids. 10 μL of each lipid extract was spotted on a silica-coated glass TLC plate. Lysyl-phosphatidylglycerol (18:1) (Avanti Polar Lipids) was used as a lipid standard. Lipids were resolved using a chloroform:methanol:water solvent system (65:25:4, v/v/v) (5). The plates were run in the solvent system for 1.5 hr and left to dry overnight, after which they were developed with iodine vapor and imaged. The plates were then left until the iodine spots disappeared completely. Ninhydrin was sprayed onto the TLC plates, which were heated on a hot plate for 5 min for the development of aminolipid spots, followed by re-imaging. Fiji (Image J) (30) was used to obtain Lys-PG intensity values. The TLC image was inverted in Image J and then quantification was performed for the Lys-PG spots. For each Lys-PG spot, a corresponding blank background spot was measured immediately next to it. Then, the corrected intensity was calculated using the following formula: corrected intensity = integrated density of Lys-PG spot - integrated density of background spot (Dataset S1).

### Murine model of bacteremia

All animal experiments were conducted under the approval of the Institutional Animal Care and Use Committee (#00316) at the University of Colorado Anschutz Medical Campus and performed using accepted veterinary standards. ∼ 7-week-old male CD-1 (Charles River) mice were challenged intravenously via tail vein injection with ∼3.5 - 5 × 10^8^ CFU of *E. faecalis* WT, *drmA*::Tn, and *mprf2*::Tn for 48 h. At 6-and-24-hour post infection, ∼10 µL blood was collected via tail bleed and at 48 hpi mice were euthanized and blood collected via cardiac puncture. At each time point, blood was serially diluted and plated on BHI or BHI supplemented with 15 µg/mL chloramphenicol agar plates to determine bacterial CFU.

### Generation of *drmA* complementation plasmid

The *E. faecalis* OG1RF *drmA* (OG1RF_11507) gene was cloned into the vector pABG5Δ*phoZ* via Gibson assembly as previously described (28) and transformed into chemically competent *E. coli* DH5*α* cells (Thermo Fisher Scientific). The plasmid sequence was confirmed at the Massachusetts General Hospital CCIB DNA Core. The sequence-confirmed plasmid, referred to as pABG5-Efs*drmA*, was electrotransformed into *E. faecalis* using previously reported methods (31, 32). Transformants of *E. faecalis* OG1RF and its derivatives were selected on kanamycin-containing agar. Transformants of *E. faecalis* V583 and its derivatives were selected on chloramphenicol-containing agar because V583 is aminoglycoside-resistant and pABG5Δ*phoZ* confers resistance to both chloramphenicol and kanamycin (27, 33). The *drmA* sequence on the complementation vector was re-confirmed directly from the transformed *E. faecalis* DAP-R strains by PCR amplification and Sanger sequencing at the Genome Center at The University of Texas at Dallas (Richardson, TX).

### Cytochrome *c* assay

The cytochrome *c* assay was performed as previously described (28). Overnight cultures of enterococcal strains were subcultured into fresh media (OD_600nm_ of 0.05), then pelleted when the OD_600nm_ reached 0.5-0.6. Cell pellets were washed twice with morpholinepropanesulfonic acid (MOPS) buffer (pH 7) and resuspended in 1 mg/mL cytochrome *c* dissolved in MOPS buffer. All samples were incubated for 10 minutes at room temperature and then pelleted by centrifugation. The absorbance of the supernatant was measured at 535 nm. The percentage of cytochrome *c* bound was calculated using the following formula: P_{cyt} = (100 – (A535/Control A535)) * 100. All assays were performed in biological triplicates for each strain.

### Gene neighborhood analysis

The Enzyme Function Initiative-Enzyme Similarity Tool (EFI-EST) (34, 35) was used to generate gene neighborhood diagrams for DUF998-encoding proteins in enterococci. The Interpro ID IPR009339, corresponding to DUF998, was used as the query to generate a sequence similarity network. Gene neighborhood diagrams were then generated to visualize genes surrounding DUF998-family protein-encoding genes.

### Heterologous expression of *E. faecium drmA* orthologs in OG1RF *drmA*::Tn

*E. faecium* 1,231,410 *drmA1* (EFTG_01541) and *drmA2* (EFTG_01549) genes were cloned into pABG5Δ*phoZ* via Gibson assembly and transformed into chemically competent *E. coli* DH5*α* cells (Thermo Fisher Scientific). The resulting plasmids were sequence-confirmed and transformed into *E. faecalis* strains as described above and are referred to as pABG5-Efm*drmA1* and pABG5-Efm*drmA2*, respectively. The transformants were verified by PCR using vector-specific primers.

### 2,4,6-Trinitrobenzene sulfonic acid (TNBS) lipid labeling

The TNBS lipid-labeling assay was performed as previously described (36–38). Overnight cultures (10 mL) were pelleted by centrifugation, washed twice and resuspended in cold HEPES (4-(2-hydroxyethyl)-1-piperazineethanesulfonic acid) buffer (pH 8.1). TNBS was added to a final concentration of 3 mM, and samples were incubated on ice in the dark for 1 hr. The reaction was terminated by the addition of lysine to a final concentration of 20 mM. The acidic Bligh Dyer lipid extraction was performed as described above, and the resulting lipid extracts were analyzed by LC-MS/MS.

### Biofilm assay by confocal microscopy

Biofilm formation was assessed as previously described (39) by modifying a previously published protocol (40). Briefly, a single colony was resuspended in 1 mL BHI and added to a sterile cover slip in a six-well plate. 4 mL BHI broth was added to each well, and the plate was incubated for 72 hours at 37℃, with the medium replaced every 24 hours. After 72 hours, the biofilms were fixed with 5% paraformaldehyde and stained with SYTO 9, green fluorescent nucleic acid stain (Invitrogen). The SYTO 9 stain was removed, and the coverslip containing the biofilm in each well was gently washed twice with 1X PBS. A drop of antifade solution (Invitrogen) was then added to a clean glass slide. A coverslip was gently removed using tweezers from the six-well plate and placed inverted on the glass slide such that the biofilm contacted the antifade solution. The biofilm was visualized via confocal microscopy (Zeiss LSM 880). The biofilm z-stack images were analyzed in ImageJ (Fiji software version 2.9.0/1.54f). Z-stack images were acquired from four different spots in the biofilm per strain. Biofilm thickness was calculated as the average of all four Z-stack readings per strain. The biofilm assay was performed in independent triplicates for each strain.

### Statistical analysis

Graphs in this study were generated by GraphPad Prism (version 9.4.1). Statistical analyses of lipid ratios, lipid abundance, TLC quantification, biofilm quantification, and cytochrome *c* assays were performed by ordinary an one-way ANOVA with multiple-comparisons testing.

### Phylogenetic analysis

The organismal taxonomic IDs from the InterPro entry IPR009339 (41) were downloaded and used to query the NCBI Taxonomy database for full lineage information (42) (acquired November 2024). These lineages were used to construct a phylogenetic tree, which was visualized using treeio and ggtree in the R programming language (43, 44). The tree was filtered to only include DUF998-containing proteins with more than 180 amino acids to ensure representation of functional DrmA variants.

## Results

### High-level DAP-resistant derivatives of *E. faecalis* V583 have drastically lower levels of the novel cationic glycolipid, Lys-Glc_2_-DAG

In a prior study, DAP-resistant (DAP-R) variants of the *E. faecalis* strain V583 were generated through up to 14 days of serial passage under escalating DAP concentrations (19). Three endpoint strains arising from independent serial passage experiments were generated, referred to as DAP-A, DAP-B, and DAP-C (Table 1). Since the DAP-R strains were generated in the 2010s (19), the DAP MIC of these strains was reconfirmed by E-test. The DAP-A and DAP-B strains have DAP MIC values in the range of 96- 128 μg/mL and DAP-C has a DAP MIC value >256 μg/mL (Table 1), consistent with their originally reported MICs (19).

Lipidomic analysis was performed for the parental V583 strain and the three DAP-R strains. The DAP-R strains had significant alterations in cationic lipid levels compared to the parental V583 strain (Figure 1). For Figure 1, the intensity values of each cationic lipid were normalized to those of the highly abundant glycolipid, Glc_2_-DAG. Notably, all DAP-R strains displayed a striking and statistically significant reduction in the Lys-Glc₂-DAG–to–Glc₂-DAG ratio. Additionally, the Lys-PG-to-Glc_2_-DAG ratios were significantly increased in the DAP-R strains DAP-A and DAP-C compared to V583. No significant alteration in the Lys-Glc-DAG-to-Glc_2_-DAG ratio was observed (Figure 1). Normalization of cationic lipids to their specific biosynthetic precursors (Figure S1A) as well as considering their absolute intensity values with no normalization (Figure S1B) also demonstrated a striking and highly significant reduction in Lys-Glc_2_-DAG levels in DAP-R strains. A trend of increased Lys-PG levels in the DAP-R strains was also observed (Figure S1A-B), albeit this was not statistically significant for all the normalization methods and comparisons.

**Figure 1.**
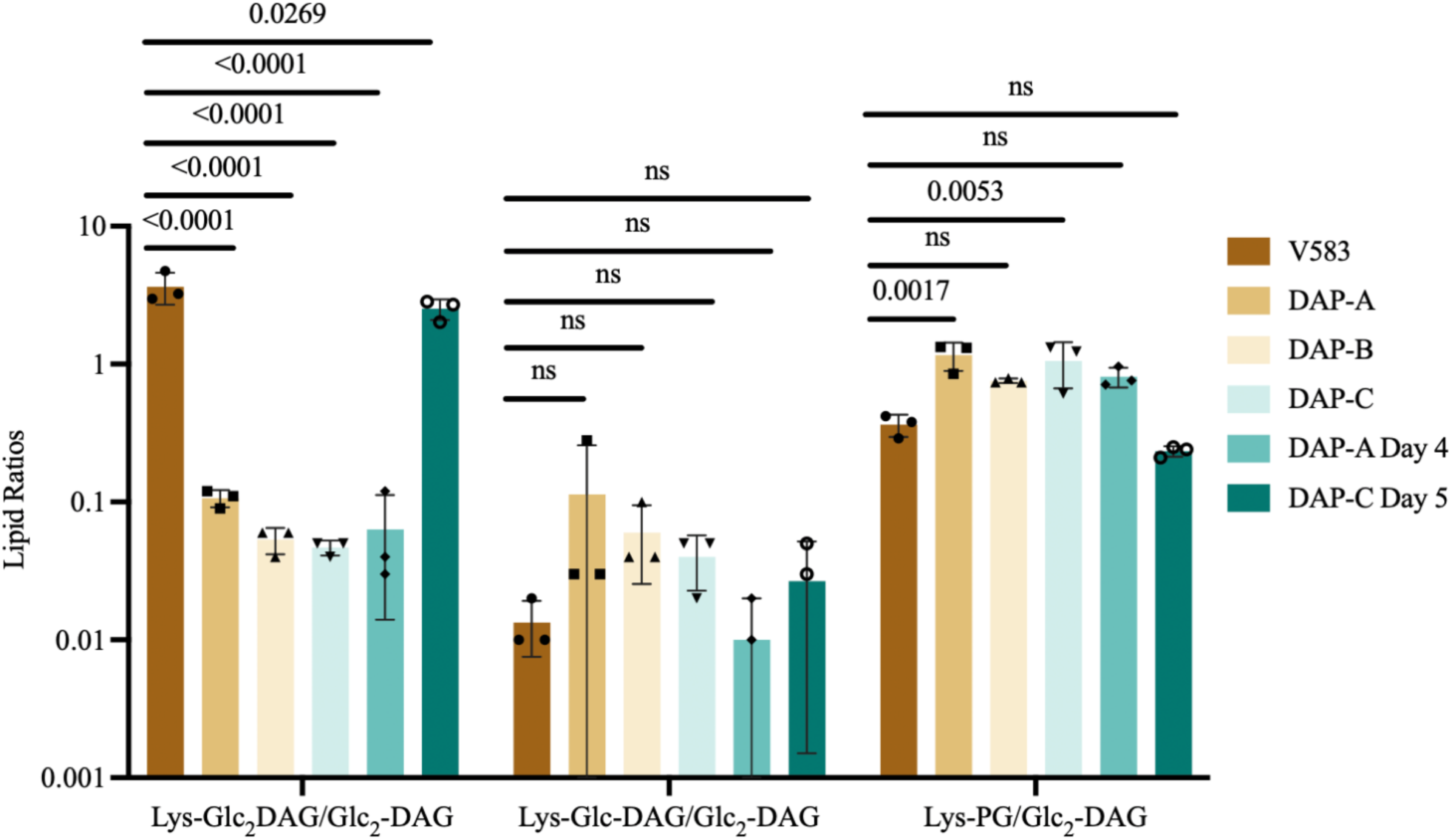
Lys-Glc_2_-DAG levels are significantly reduced in DAP-R strains with *drmA* mutations compared to those without. Intensity values for each lysine-modified lipid were normalized to those of the glycolipid Glc_2_-DAG (n=3 independent replicates). Bars represent the mean lipid ratio across replicates ± standard deviation. Significance was assessed using ordinary one-way ANOVA with Dunnett’s multiple comparisons test.

Additionally, the DAP-A and DAP-C strains have altered acyl chain distributions compared to each other, and the parental V583 and DAP-B strains. Figure S2 shows representative free fatty acid profiles (panel A), and Glc_2_-DAG and PG acyl chain distributions (panels B and C, respectively), for these strains. The DAP-B profiles are largely superimposable over the parental V583 profiles for all panels. For DAP-A, the free fatty acids and acyl side chains are shifted towards shorter chains, with Glc_2_-DAG (14:0/16:1) and the PG species (16:0/14:1) and (16:1/17:1) being unique to DAP-A (Figure S2). For DAP-C, the free fatty acid profile and PG species predominantly consist of saturated chains (Figure S2).

### Lys-Glc_2_-DAG reduction in DAP-R strains co-occurs with loss-of-function mutations in *drmA*

These results led us to determine whether the drastic reduction in Lys-Glc_2_-DAG levels was linked to specific gene(s) in the DAP-R strains, which had previously been analyzed by genome sequencing (Table 1). All endpoint strains from the three passage experiments harbored mutations in at least two genes: EF0631 (*cls1*), encoding cardiolipin synthetase, and a gene of unknown function, encoded by V583 locus EF1797, previously named *drmA* (19, 20). The *cls1* allele and its role in enterococcal DAP resistance has been investigated (8, 19, 20, 45–47). In contrast, the role of *drmA* has not been determined, although presumptive loss-of-function mutations in this gene have been associated with enterococcal DAP resistance in multiple studies (8, 19–23) (Table S1) and with increased biofilm formation in DAP-R strains (20).

We hypothesized that mutation of *drmA* was associated with the lower Lys-Glc_2_-DAG levels observed in the DAP-R strains. Strains isolated earlier in the DAP serial passage experiments harbored fewer mutations and exhibited lower DAP MICs than the endpoint DAP-R strains (Table 1). Two strains from mid-points in the serial passage experiments were selected for lipidome analysis: DAP-A Day 4, which had mutations in *cls1* and *drmA*, and DAP-C Day 5, which had mutations in *cls1* and EF1753, encoding LiaX. The DAP-A Day 4 strain has dramatically reduced Lys-Glc_2_-DAG levels compared to the wild-type strain, whereas the DAP-C Day 5 strain does not (Figure 1, S1). Thus, all DAP-R strains with *drmA* mutations (DAP-A, DAP-B, DAP-C, and DAP-A Day 4) had significantly lower Lys-Glc_2_-DAG levels than strains lacking *drmA* mutations (V583 parent and DAP-C Day 5), consistent with a role for *drmA* in regulation of Lys-Glc_2_-DAG levels.

### *drmA* modulates Lys-Glc_2_-DAG levels

To determine whether *drmA* regulates Lys-Glc_2_-DAG levels, we analyzed its ortholog (OG1RF_11507) in *E. faecalis* OG1RF using a mutant (referred to as *drmA*::Tn hereafter) from a previously reported transposon mutant library (26). We observed a significant reduction in Lys-Glc_2_-DAG levels in the *drmA*::Tn strain compared to the wild-type strain in both normalized and unnormalized lipidomic data (Figure 2A and Figure S3). This defect could be complemented by *drmA* expression from the plasmid vector pABG5 (Figure 2A and Figure S3). Together, these data conclusively link *drmA* to Lys-Glc_2_-DAG levels in *E. faecalis*, demonstrating that loss of *drmA* function leads to significantly decreased Lys-Glc_2_-DAG levels.

**Figure 2.**
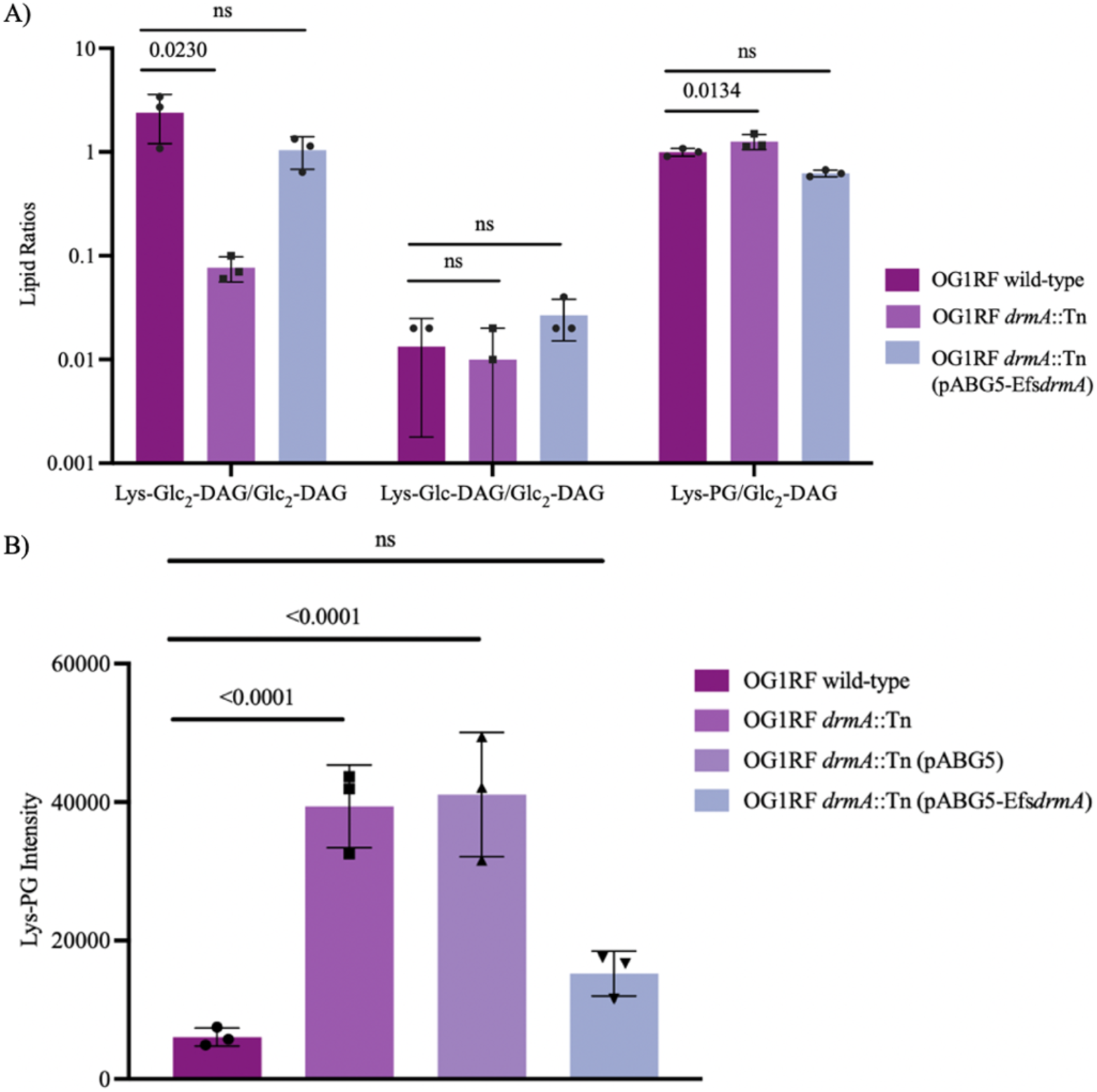
Lys-Glc_2_-DAG is significantly reduced in *drmA*::Tn with a compensatory increase in Lys-PG levels. A) Lys-Glc_2_-DAG levels are significantly reduced in *drmA*::Tn compared to OG1RF wild-type strain. Intensity values for each lysine-modified lipid were normalized to those of the glycolipid Glc_2_-DAG (n=3 independent replicates). B) Bar graph indicates TLC quantification of Lys-PG intensity across different strains. The *p*-value of the statistically significant changes are labeled in the graph. *In A) and B) bars represent the mean lipid ratio ± standard deviation and mean Lys-PG intensity ± standard deviation. Significance was assessed using ordinary one-way ANOVA for Dunnett’s multiple comparisons test.

### Lys-PG is increased in the *E. faecalis* OG1RF *drmA*::Tn

The data in Figure 2 suggest a small but significant increase in Lys-PG levels in the *drmA*::Tn mutant, which was complemented back to wild-type levels. However, this increase was not observed when using an alternative normalization method or in unnormalized data (Figure S3). We also observed increased Lys-PG levels in the *E. faecalis* DAP-R strains with *drmA* mutations, albeit not statistically significant for all normalization methods and comparisons (Figure 1 and Figure S1A-B). To further assess the impact of loss of *drmA* function on Lys-PG production, we performed additional lipid analysis using *E. faecalis* OG1RF and its derivatives. Lys-PG levels were assessed using one-dimensional thin-layer chromatography (1D-TLC) for the overnight cultures of *E. faecalis* OG1RF wild-type, *drmA::*Tn, *drmA*::Tn (pABG5) and *drmA*::Tn (pABG5-Efs*drmA*). No significant differences in growth yield were observed among these strains (OD_600nm_ values; one-way ANOVA for multiple comparisons, p > 0.05). The 1D-TLC results indicate that *drmA::*Tn, *drmA*::Tn pABG5 have larger Lys-PG spots in all three replicates under both iodine and ninhydrin staining conditions (Figure S4D and Figure S4A-B). In contrast, the wild-type strain and the *drmA* complementation strain have smaller Lys-PG spots consistently across the replicates (Figure S4D and Figure S4A-B). The TLC quantification data for Lys-PG indicate that Lys-PG levels are higher in *drmA::*Tn (p-value <0.0001) and *drmA*::Tn pABG5 (p-value <0.0001) relative to the OG1RF wild-type and *drmA* complementation strain (Figure 2C, Figure S4C, and Dataset S1). These data indicate a potential seesaw effect with Lys-Glc_2_-DAG and Lys-PG; as Lys-Glc_2_-DAG levels decrease, Lys-PG increases.

Notably, all strains shown in Fig. 2 presented DAP MICs of 1-1.5 μg/mL (Dataset S1). Thus, the loss of *drmA* function and the resulting changes in Lys-Glc_2_-DAG and Lys-PG levels did not alter DAP susceptibility in the OG1RF background. To assess if *drmA* contributes to bloodstream survival, mice were challenged with OG1RF, *drmA::*Tn, and *mprf2::*Tn and blood collected at 6, 24, and 48 hour post infection (hpi). Loss of *drmA* did not affect blood survival compared to OG1RF, whereas the loss of *mprf2* resulted in decreased (non-significant) CFU recovered in the bloodstream after 48 hpi, as has previously been observed (7) (Figure S5). Hence, *drmA* inactivation in OG1RF does not confer significant protection from daptomycin nor from the murine immune system.

### Restoration of *drmA* function reverses DAP MIC in high-level DAP-R *E. faecalis* V583

We reasoned that if *drmA* loss-of-function contributed to DAP resistance, then complementing the DAP-R strains with *drmA* would result in lower daptomycin MIC. Indeed, we observed a significant decrease in DAP MIC when OG1RF *drmA* was expressed *in trans* in these strains (Figure 3A), but not full reversion to DAP susceptibility (per clinical breakpoints (48)). As expected, *drmA* complementation restored Lys-Glc_2_-DAG levels (red peak) to the DAP-R strains (Figure 3B). Note that the two peaks observed on the chromatograms for Lys-Glc_2_-DAG correspond to lysine linkage to either the first or second glucose of Glc_2_-DAG (confirmed by MS/MS) (Figure S6). Our results are consistent with *cls1* mutation being a “first step” towards high-level DAP resistance in the V583 background, after which *drmA* mutation contributes to progression of resistance towards higher MICs. Hence, *drmA* loss-of-function played a role in the progression to high-level DAP-R in V583. Considering our OG1RF *drmA*::Tn results, in which *drmA* loss-of-function did not impact DAP MIC (Figure 2 and Dataset S1), our results suggest that alterations in cationic lipids alone are not sufficient to confer DAP resistance. Rather, we hypothesize that inactivation of *drmA* must occur on the background of prior adaptive mutations (for e.g., in *cls1*) to confer elevated DAP MIC.

**Figure 3.**
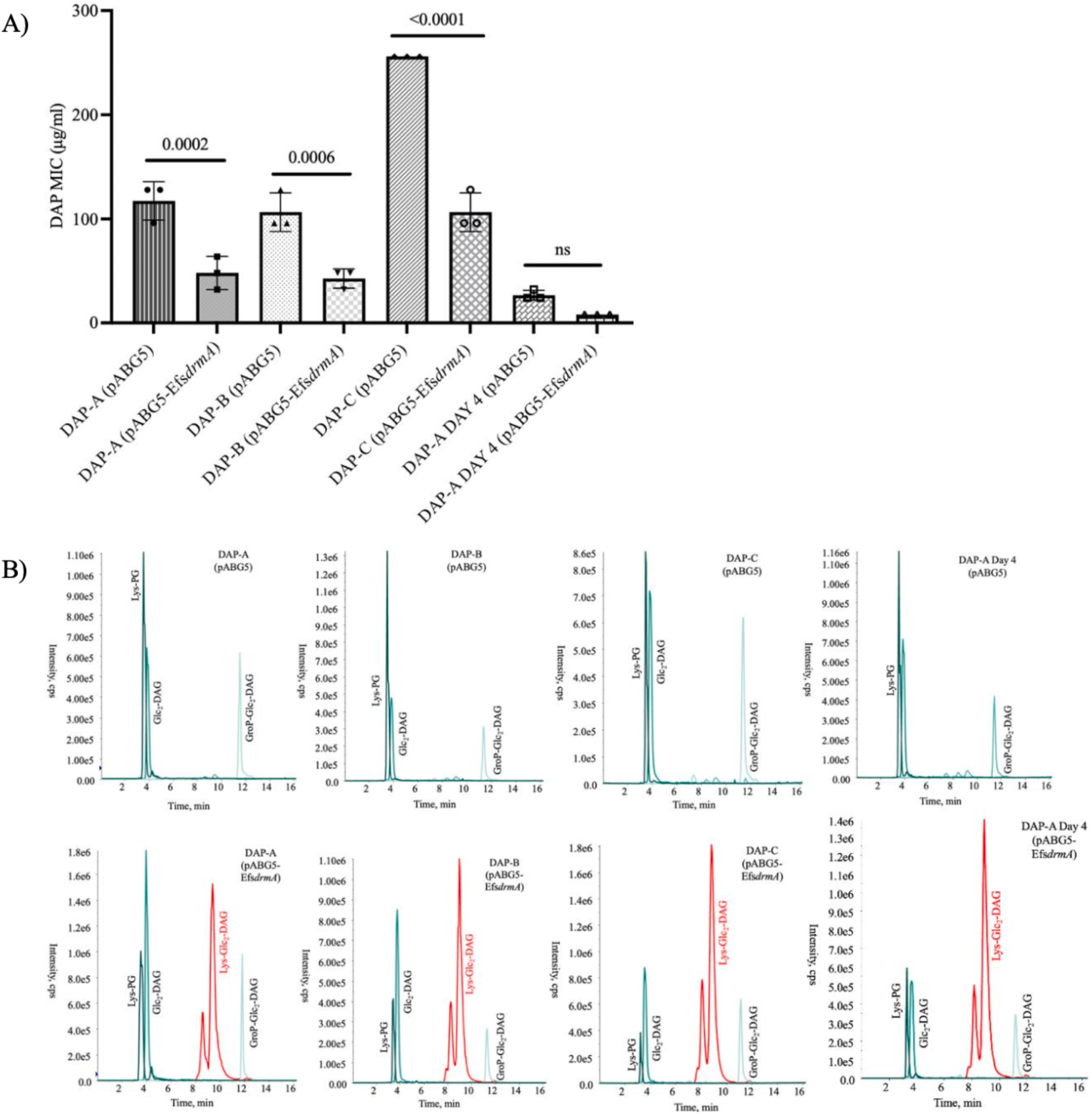
Complementation with *drmA* reverses DAP MIC in DAP-R *drmA* mutants and restores Lys-Glc_2_-DAG levels. A) The graph represents DAP MIC (µg/mL) of DAP-R empty vector strains and their corresponding *drmA* complemented DAP-R strains for n=3 independent replicates. The bars represent mean with standard deviation. Significance was assessed using ordinary one-way ANOVA for Dunnett’s multiple comparisons test. (B) The X-axis in the ion chromatograms indicates retention time in minutes and the Y-axis indicates the intensity. In the DAP-R empty vector strains Lys-PG is seen at retention time 3-4 minutes on the X-axis, Glc_2_-DAG is seen at retention time 4-5 minutes on the X-axis, GroP-Glc_2_-DAG is seen at retention time 11-12 minutes on the X-axis. In the *drmA* complemented DAP-R strains Lys-Glc_2_-DAG is seen at retention time 8-10 minutes on the X-axis and the other cationic lipids are seen at the same retention time as mentioned above. Abbreviations: Glc_2_-DAG, –diglucosyl-diacylglycerol; Lys-Glc_2_-DAG, -lysyl-diglucosyl-diacylglycerol, GroP-Glc_2_-DAG, -glycerophospho-diglucosyl-diacylglycerol.

### One of the two *E. faecium drmA* orthologs complements Lys-Glc_2_-DAG levels in *E. faecalis drmA*::Tn, but both orthologs complement its biofilm phenotype

*E. faecalis* encodes a single copy of *drmA* but *E. faecium* encodes two orthologs of *drmA* (Figure 4A; (19)). The enterococcal DrmA protein structures indicate high structural similarity but limited percent amino acid identity (Figure 4B and Dataset S1). We hypothesized that the *E. faecium drmA* orthologs might also impact Lys-Glc_2_-DAG levels. We found that expression of *E. faecium drmA1* from pABG5-Efm*drmA1* complemented Lys-Glc_2_-DAG levels in *drmA*::Tn, but expression of *E. faecium drmA2* from pABG5-Efm*drmA2* did not (Figure 4C).

**Figure 4.**
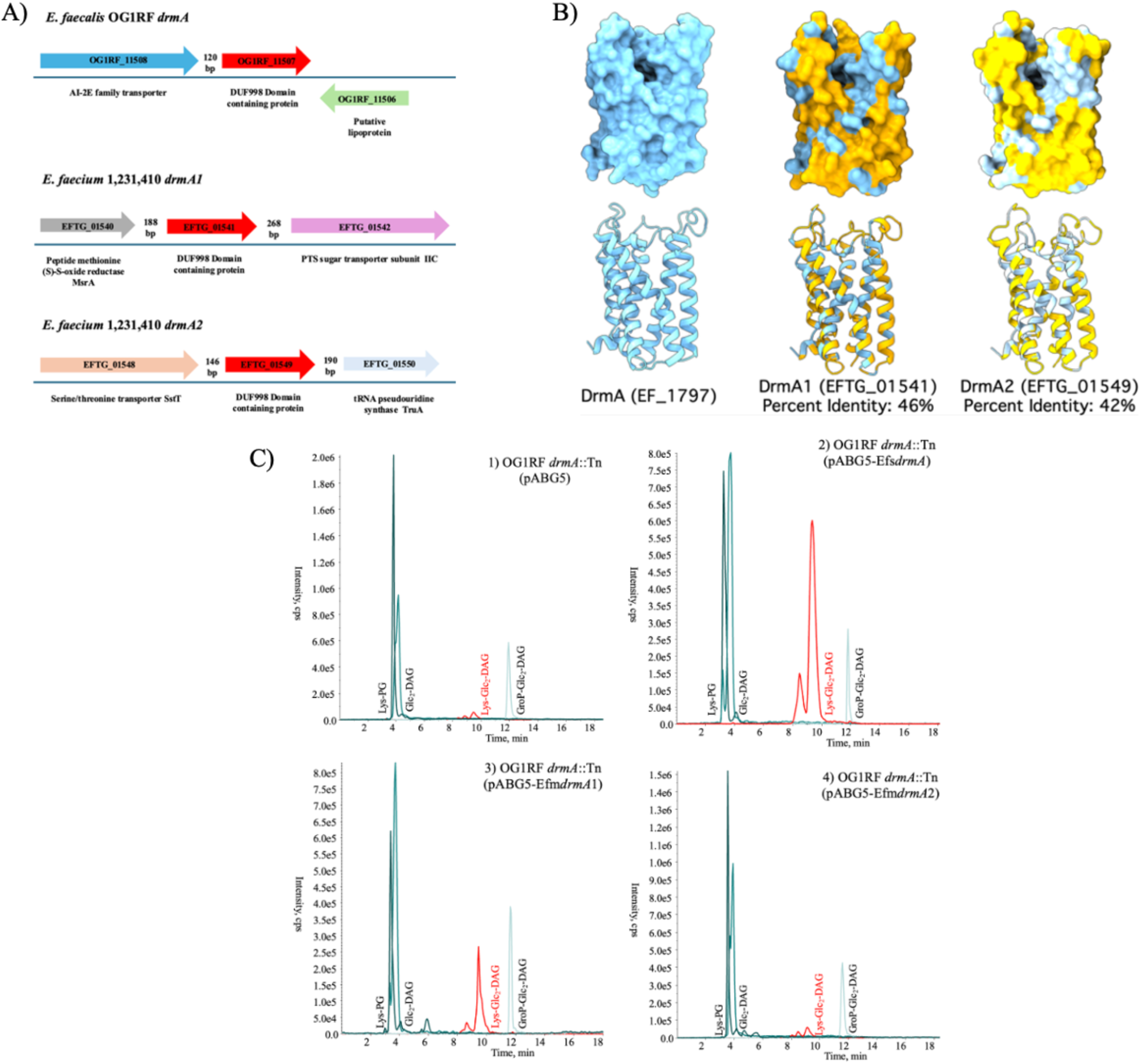
One of the two *E. faecium drmA* orthologs can complement Lys-Glc_2_-DAG in *E. faecalis drmA*::Tn. A) *drmA* (red arrows) gene organization in *E. faecalis* OG1RF and *E. faecium* 1,231,410 genomes. (B) The predicted structures of enterococcal DrmA in *E. faecalis* V583 and *E. faecium* 1,231,410, with percent amino acid identity of the *E. faecium* DrmA proteins to the *E. faecalis* DrmA shown. (C) Ion chromatograms showing the Glc_2_-DAG and Lys-Glc_2_-DAG intensities in: OG1RF *drmA*::Tn with empty vector pABG5 (panel 1), complementation strain OG1RF *drmA*::Tn (pABG5-Efs*drmA*), heterologous expression of *E. faecium drmA1* (EFTG_01541) in OG1RF *drmA*::Tn (Panel 3) and *E. faecium drmA2* (EFTG_01549) in OG1RF *drmA*::Tn (Panel 4). Abbreviations: Glc_2_-DAG, - diglucosyl-diacylglycerol; Lys-Glc_2_-DAG, - lysyl-diglucosyl-diacylglycerol

Prior work found that mutations in *drmA* contribute to the aggressive biofilm formation phenotype observed for laboratory-evolved DAP-R *E. faecalis* (20). However, the individual role of *drmA* in biofilm formation has not been assessed. We cultured *E. faecalis* biofilms on plastic coverslips for 72 hours, followed by staining and visualization by confocal microscopy (Figure 5 and Figure S7). We observed greatly enhanced biofilm thickness for *drmA*::Tn, which was complemented back to wild-type levels by *in trans* expression of *E. faecalis drmA1*, *E. faecium drmA1* or *E. faecium drmA2* (Figure 5). It is unclear whether Lys-Glc_2_-DAG plays a direct role in *drmA*-altered biofilm thickness. Recent work in *S. aureus* revealed that Lys-PG aids in biofilm formation by enhancing cell-to-cell interaction and thereby reducing electrostatic repulsion among negatively charged bacterial surfaces (49). It is possible that the increase in Lys-PG that we observed for *drmA*::Tn is the actual mediator of increased biofilm formation. However, expression of *E. faecium drmA1* complemented Lys-Glc_2_-DAG in *drmA*::Tn, but expression of *E. faecium drmA2* did not (Figure 4C), perhaps suggesting that the roles of *drmA* in Lys-Glc_2_-DAG regulation and in biofilm formation are separatable. The interplay of Lys-Glc_2_-DAG and Lys-PG synthesis and their roles in biofilm formation in *E. faecalis* remain to be investigated.

**Figure 5.**
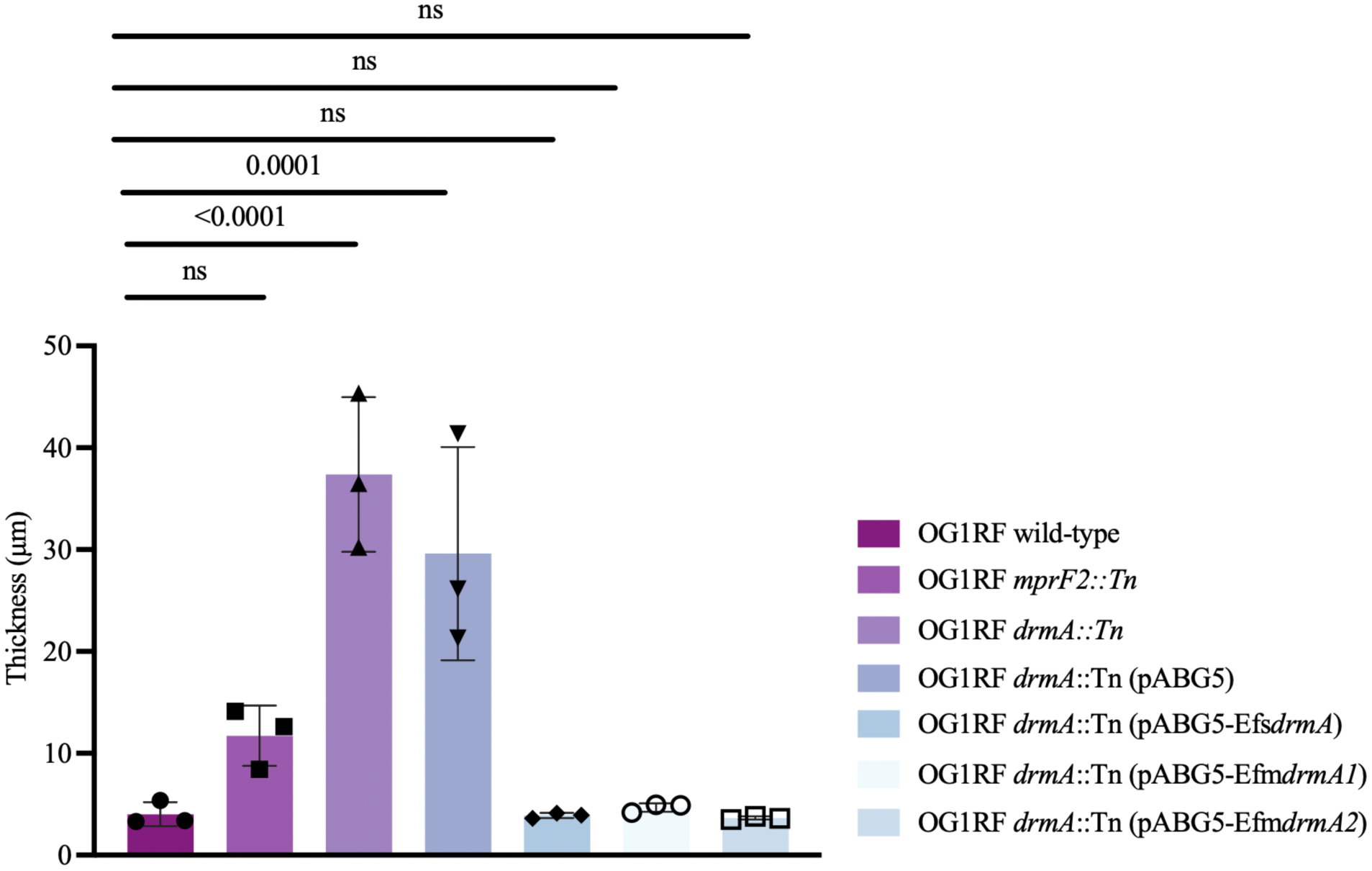
*E. faecalis* OG1RF *drmA*::Tn forms significantly thicker biofilms, which can be complemented by both *E. faecium drmA* alleles. The graph indicates the biofilm formation of the strains *E. faecalis* OG1RF wild-type, *mprF2*::Tn, *drmA*::Tn, OG1RF*drmA*::Tn (pABG5), OG1RF*drmA*::Tn (pABG5Efs*drmA*), OG1RF*drmA*::Tn (pABG5-Efm*drmA1*), and OG1RF*drmA*::Tn (pABG5-Efm*drmA2*) (n=3 independent replicates). Bars represent the mean ± standard deviation. Significance was assessed using ordinary one-way ANOVA for Dunnett’s multiple comparisons test.

### Lys-Glc_2_-DAG is present on the external surface of the *E. faecalis* cell membrane

MprF possesses a flippase domain that translocates aminoacylated lipids from the inner leaflet, where they are synthesized, to the outer membrane leaflet, where they face the external environment. The MprF flippase domain function has been primarily studied in *Staphylococcus aureus* and *Pseudomonas aeruginosa* (50, 51); however, these organisms do not synthesize cationic glycolipids (9). It is unknown whether cationic glycolipids like Lys-Glc_2_-DAG can be flipped by MprF or other flippases as the membrane leaflet localization of Lys-Glc_2_-DAG has not been investigated.

The membrane-impermeable reagent 2,4,6-trinitrobenzenesulfonic acid (TNBS) has been used to analyze aminophospholipid asymmetry (37, 38). TNBS has been used to label the lipid phosphatidylethanolamine (PE) in *Bacillus megaterium* (37) and in *Escherichia coli* (38), confirming its asymmetric distribution in the membranes of these bacteria. TNBS cannot cross the membrane due to its water solubility and net negative charge and hence, it only derivatizes aminolipids in the outer leaflet (37). It reacts with primary amines on lipids to form trinitrophenylated lipid products that can be detected by LC-MS/MS (38). As a control, we treated the Gram-positive model bacterium *Bacillus subtilis* with TNBS, confirming that PE, a major phospholipid synthesized by this organism, is labeled with TNBS, as expected (Figure 6A). TNBS also labeled Lys-PG in *B. subtilis* (Figure 6A), which encodes *mprF* (52–54). Together, these results confirm that TNBS can be used to analyze the outer-leaflet localization of multiple aminolipids in bacterial membranes.

**Figure 6.**
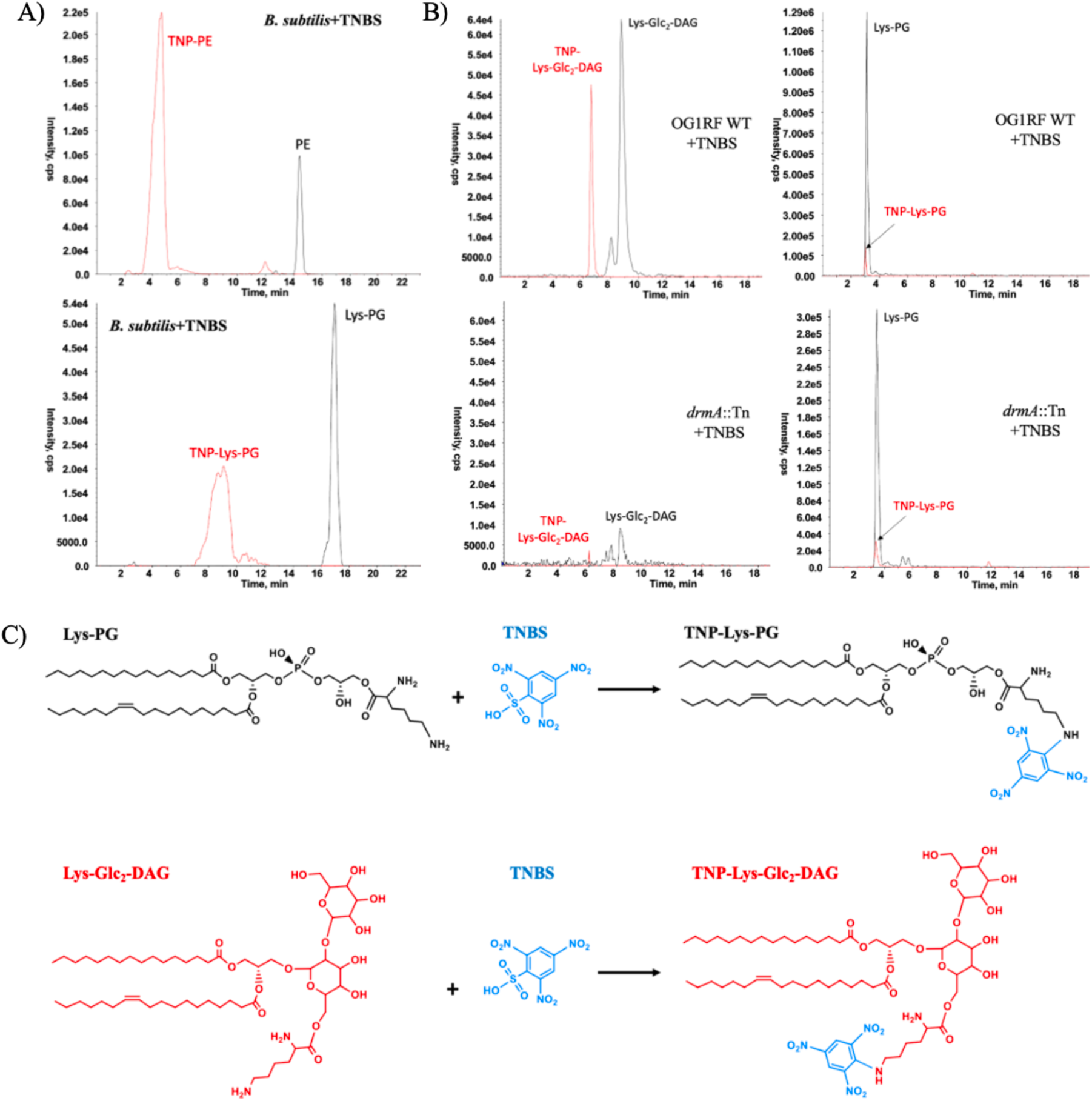
TNBS supports Lys-Glc_2_-DAG presence in the outer leaflet of the enterococcal membrane. (A) and (B) The X-axis of the ion chromatogram represents retention time in minutes, and the Y-axis represents intensity. (A) In *Bacillus subtilis,* the TNBS-labeled phosphatidylethanolamine (PE) product TNP-PE is seen at retention time 3-5 minutes and the TNBS-labeled Lys-PG product TNP-Lys-PG is seen at retention time 7-10 minutes in the chromatogram. (B) In OG1RF wild-type, unlabeled Lys-Glc_2_-DAG is seen at retention time 8-10 minutes and TNBS–labeled Lys-Glc_2_-DAG is seen at 6-7 minutes in the chromatogram. In *drmA*::Tn, trace amount of Lys-Glc_2_-DAG is seen at retention time 8-10 minutes, but its TNBS-labeled product is absent. In OG1RF wild-type and in *drmA*::Tn, the unlabeled Lys-PG is seen at retention time 3-4 minutes and TNBS–labeled Lys-PG is seen at retention time 3 minutes in the chromatogram. (C) The chemical reaction depicts that TNBS reacts with primary amines in *E. faecalis* lipids to form trinitrophenylated (TNP) lipid products.

**Figure 7.**
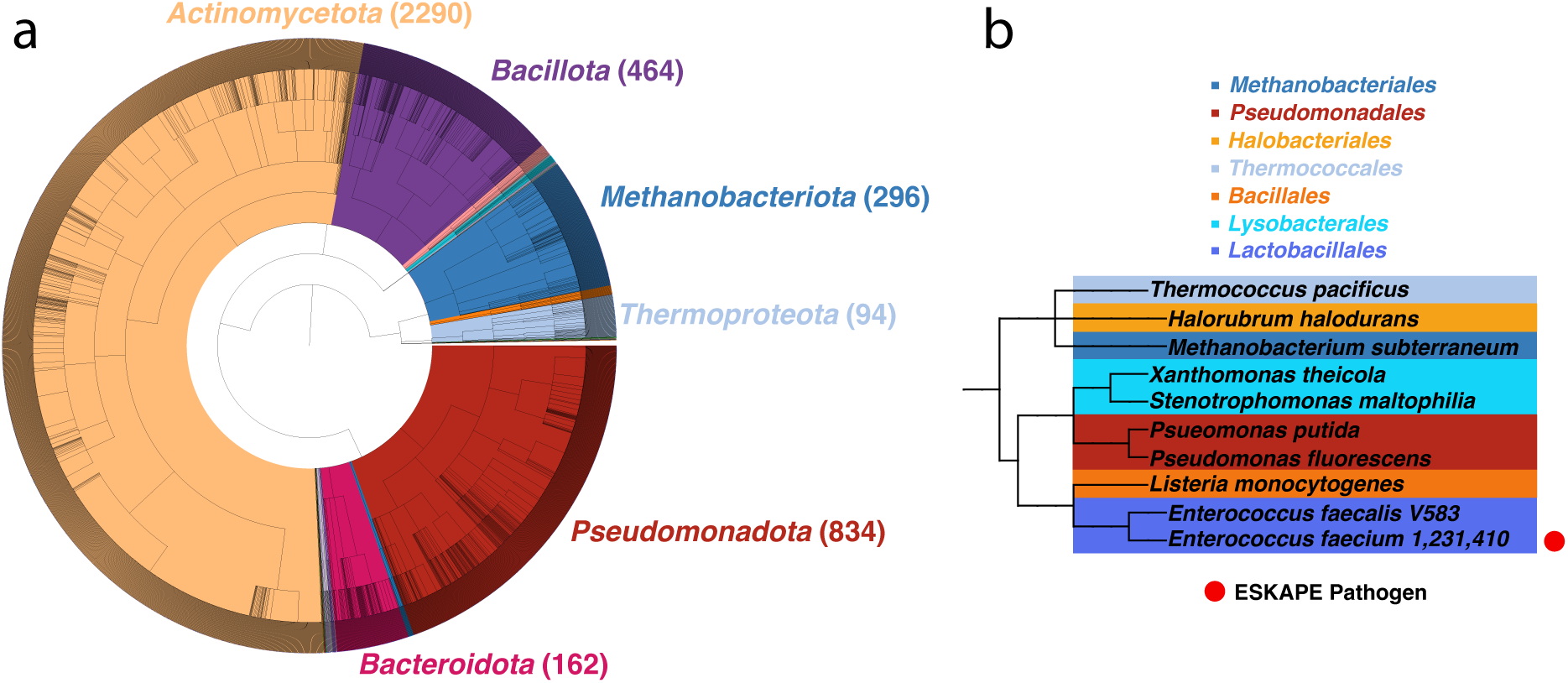
Phylogenetic analysis of the DUF998 family (PF06197). (a) Phylogenetic tree built using full taxonomic lineage from the Interpro dataset for DUF998. (b) Small tree of species mentioned in text, with ESKAPE pathogens containing a DUF998 domain protein highlighted. Legend contains the specific Phylum the listed species belong to.

TNBS labeling of *E. faecalis* OG1RF yielded robust labeling of Lys-Glc_2_-DAG, indicating that this lipid is flipped to the outer leaflet in *E. faecalis* OG1RF (Figure 6B). In comparison, TNBS-labeled and unlabeled Lys-Glc_2_-DAG are detected at trace levels in the *drmA*::Tn mutant (Figure 6B). TNBS-labeled Lys-PG was detected in both wild-type and *drmA*::Tn strains (Figure 6B). Chemical reactions for the TNBS modification of enterococcal lysine-modified lipids are shown in Figure 6C. As expected, no TNBS-labeled or -unlabeled cationic lipids were detected in the *mprF2*::Tn mutant (negative data not shown). *E. faecalis* does not synthesize PE, nor does it encode known pathways for PE synthesis, and this lipid was not detected.

Given this evidence that Lys-Glc_2_-DAG is flipped to the outer membrane leaflet of *E. faecalis*, we reasoned that the differences in Lys-Glc_2_-DAG levels in the DAP-R strains and in the *drmA*::Tn mutant would result in substantial alterations to cell surface charge. However, cytochrome c binding assays, which are a measure of relative differences in cell surface charge, revealed no significant differences among *E. faecalis* V583 and its DAP-A and DAP-C derivatives, despite substantial differences in lipid profiles and DAP MICs among these strains (Figure S8). Moreover, no significant differences were observed among *E. faecalis* OG1RF and its *drmA*::Tn and *mprF2*::Tn derivatives, despite substantial differences in lipid profiles (Figure 2 and our previously published *mprF2*::Tn lipidome data (9)) (Figure S8). This is in contrast with another Gram-positive pathogen, *Streptococcus agalactiae,* in which the loss of lysine-modified lipids (Lys-Glc-DAG and Lys-PG) in an *mprF* deletion strain was associated with a significantly reduced net positive charge as assessed by cytochrome c binding (55). Our data suggest that an increase in Lys-PG in *drmA*::Tn compensates for the loss of Lys-Glc_2_-DAG to balance overall enterococcal surface charge, but this requires additional investigation.

### Phylogenetic analyses reveal widespread distribution of DUF998/DrmA family proteins in prokaryotes

To assess how widespread DrmA is across the tree of life, we performed a phylogenetic analysis of the protein. DrmA is a member of the Domain of Uncharacterized Function 998 family (PF06197) from the Pfam database of protein domains (56). While this domain is uncharacterized, it belongs to the Pfam clan for Frag1-like proteins, which is a broad family found predominantly in eukaryotes but also present in some bacterial and archaeal species. The Frag1-like proteins are transmembrane proteins with a six–α-helical fold and have been implicated in the regulation of lipid metabolism and cell signaling in eukaryotes (57, 58). Consistent with this classification, the predicted structure of *E. faecalis* DrmA from the AlphaFold database (59) also exhibits a six–α-helical fold (Figure 8A). To our knowledge, neither Frag1-like proteins nor DUF998 proteins have crystal structures available.

**Figure 8.**
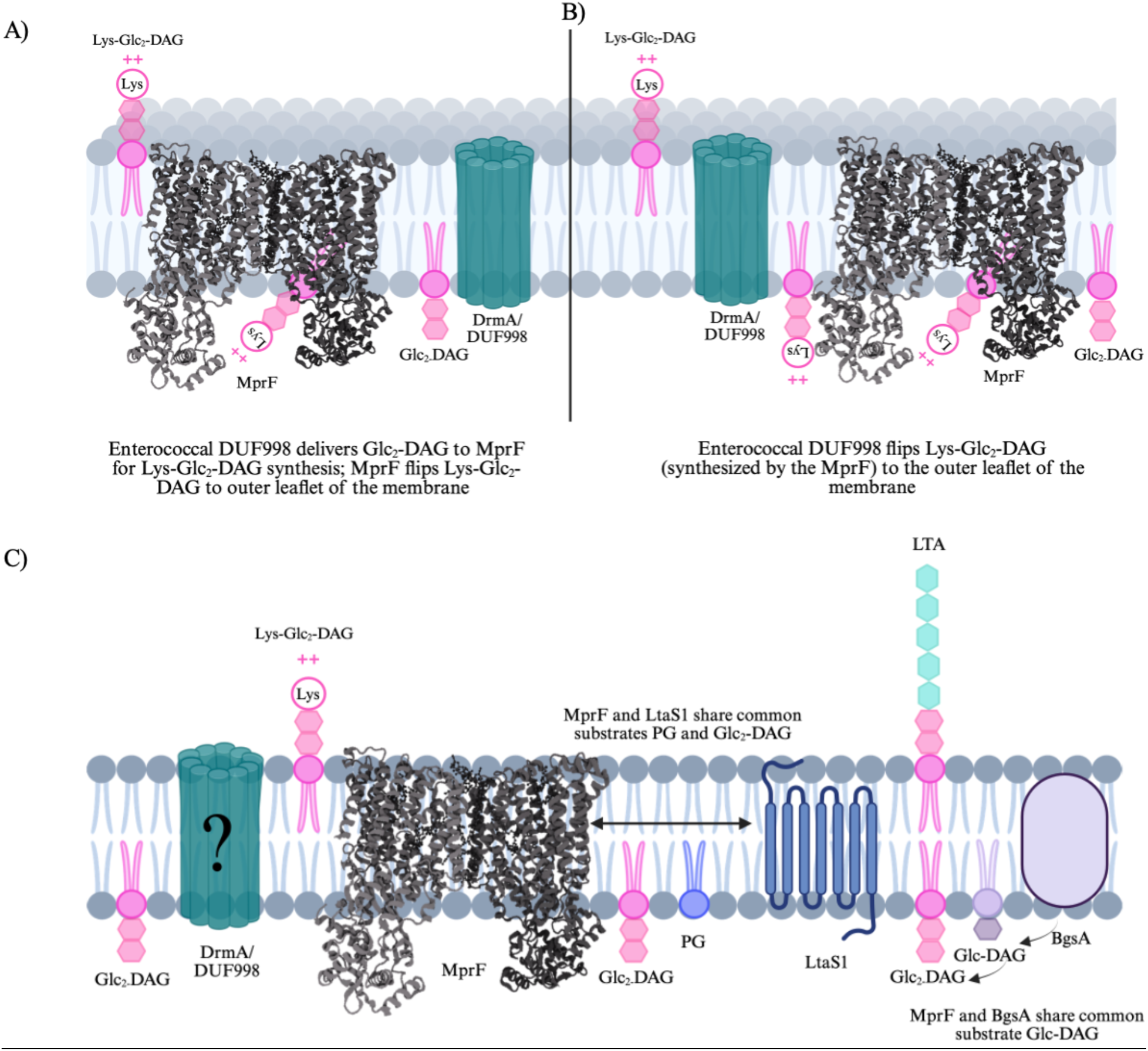
Proposed Model for the function of DrmA and interconnections among pathways for lysinylated lipids and Type I lipoteichoic acid (LTA). (A) DrmA might have substrate specificity to Glc_2_-DAG and interact with the MprF2 synthase domain by delivering Glc_2_-DAG to synthesize Lys-Glc_2_-DAG. Then, the MprF flippase domain most likely flips Lys-Glc_2_-DAG to the outer leaflet of the membrane. (B) Alternatively, DrmA could have specificity to Lys-Glc_2_-DAG and could be the designated flippase for Lys-Glc_2_-DAG. Lys-Glc_2_-DAG might have a specific flippase due to its relatively distinct and bulky structure compared to the other enterococcal lysine lipids Lys-Glc-DAG and Lys-PG. C) MprF in *E. faecalis* utilizes PG and Glc_2_-DAG as substrates, both of which are also used for Type I LTA synthesis by the enzyme LtaS1. MprF additionally utilizes Glc-DAG as a substrate, which is also used by the Glc_2_-DAG synthesis enzyme BgsA.

We found that across the ∼11,000 sequences found in the Uniprot database for PF06197, the dominant hosts of this protein domain are bacteria and archaea (Figure 8B), but with uneven distribution among and within clades. For example, the domain is present in some but not all enterococci. For instance, it is found in *E. faecalis* and *E. faecium* but not in *E. gallinarum*. Among Gram-positive pathogens, the domain is present in *Listeria monocytogenes* (encoded by locus *lmo0296* in the model strain EGD-e), but not in *S. aureus* or major streptococcal pathogens, including *S. pneumoniae* or *S. pyogenes*. Notably, *S. agalactiae* does not encode this domain, yet *S. agalactiae* synthesizes Lys-Glc-DAG (28). Thus, the distribution of this domain is not strictly correlated with cationic glycolipid synthesis. Among major Gram-negative pathogens, the domain is present in *Stenotrophomonas maltophilia* and *Xanthomonas* sp., but is absent from *E. coli*, *Klebsiella*, *Acinetobacter*, and *Pseudomonas aeruginosa* (albeit present in *P. putida* and encoded by locus PP_5607 in the model strain KT2440). Beyond pathogens, the domain has a broad distribution among more environmentally-associated taxa, including among the *Actinomycetes* and the archaeal genera such as *Thermococcus*, *Methanobacterium*, and *Halorubrum*. Overall, we conclude that DrmA is widespread among prokaryotes but is not a core protein, and its distribution does not correlate with specific membrane lipid compositions, which vary widely across the taxa it is present in (24, 60). Investigation of DrmA-like proteins in microbes beyond enterococci is therefore warranted to further understand their functions and contributions to membrane lipid composition and biofilm formation.

## Discussion

Motivated by our recent discovery of cationic glycolipids in enterococci (9), we revisited a prior collection of high-level DAP-R *E. faecalis* strains (19) to investigate whether these novel lipids contribute to DAP resistance emergence. Indeed, we found that levels of the cationic glycolipid Lys-Glc_2_-DAG were dramatically lower in the DAP-R strains relative to the parental wild-type strain. We further established that inactivating mutations in *drmA*, encoding a DUF998 family protein of unknown function, conferred this dramatic reduction in Lys-Glc_2_-DAG levels. Moreover, restoration of *drmA* to the DAP-R strains lowered their DAP MICs, reversing their trajectory to high-level resistance, demonstrating that *drmA* loss-of-function is an important contributor to high-level DAP resistance. Crucially, the effect of *drmA* loss-of-function on DAP MIC appears to be epistatic with preceding mutations on the path to high-level resistance (most likely in the cardiolipin synthase-encoding *cls1*, but this remains to be investigated), as inactivation of *drmA* in the non-DAP-R strain *E. faecalis* OG1RF did not alter DAP MIC.

Our study has established that DrmA is significantly associated with Lys-Glc_2_-DAG levels, and with Lys-PG levels, albeit to a lower magnitude. What is the molecular mechanism of DrmA function? Assuming direct interactions, DrmA could interact with either the substrates (Glc_2_-DAG or lysyl-tRNA) or product (Lys-Glc_2_-DAG) of the Lys-Glc_2_-DAG synthesis reaction. We note that MprF is required for the synthesis of Lys-Glc_2_-DAG in *E. faecalis* (9) and is likely sufficient, given that our experiments in this study found that *drmA* mutants continue to synthesize a low level of Lys-Glc_2_-DAG. We propose that DrmA interacts with Glc_2_-DAG, Lys-Glc_2_-DAG, or both. In one plausible model for DrmA function, DrmA binds Glc_2_-DAG and delivers it to MprF, thereby facilitating Lys-Glc_2_-DAG synthesis. In the absence of DrmA, Glc_2_-DAG is either poorly available or poorly recruited to MprF, resulting in low levels of Lys-Glc_2_-DAG synthesis and increased Lys-PG synthesis. The distribution of MprF in the membrane relative to its lipid substrates (and to DrmA) is unknown, as is true for most membrane proteins in enterococci; this would be valuable information for the field. In a second potential model for DrmA function, MprF synthesizes cationic lipids, but cannot flip Lys-Glc_2_-DAG to the outer leaflet; rather, DrmA is the Lys-Glc_2_-DAG flippase. In the absence of DrmA, Lys-Glc_2_-DAG accumulates in the inner leaflet and/or in the MprF enzyme itself, resulting in feedback inhibition of MprF and decreased synthesis of Lys-Glc_2_-DAG. For either of these two models, an important next step will be to determine whether DrmA is capable of binding the relevant lipids. Our data suggests a broader role of DrmA, in that DrmA is part of a lipid regulatory network. We hypothesize that DrmA and MprF are tightly coupled during enterococcal adaptation to DAP and that they operate in a synchronized network to regulate cell membrane homeostasis.

Another key question is how *drmA* loss of function mechanistically contributes to DAP resistance progression. Our work in the V583 and OG1RF strain backgrounds suggests that loss of *drmA* function alone, and its concomitant impact on cationic lipid levels, is not sufficient to confer significant DAP MIC changes; rather, its effect is epistatic with other mutations. Further genetic work will be required to fully support this model, for example, introducing *cls1* and *drmA* mutations simultaneously into OG1RF, coupled with DAP MIC testing and lipidomic analyses to link lipid alterations with MIC. A recent study by Rashid et al. elucidated the various roles MprF plays in the intracellular network that greatly influence membrane plasticity (5). Notably, MprF shares common substrates (PG and Glc_2_-DAG) with the Type I LTA biosynthesis. Recent work shows that LtaS1 in *E. faecalis* is responsible for LTA synthesis and also suggests that MprF products may be recycled into LtaS1-directed LTA synthesis (61). Hence, strains harboring mutations in *drmA* resulting in significant loss of Lys-Glc_2_-DAG levels might have altered LTA levels or composition as a result of these interconnected pathways of substrate utilization. For example, if Lys-Glc_2_-DAG synthesis competes with LTA synthesis for Glc_2_-DAG substrate, then reduced Lys-Glc_2_-DAG levels should result in more Glc_2_-DAG availability for LtaS1. Future studies seeking to understand the role of DrmA and cationic lipids in high-level DAP-R should include analysis of LTA composition, length, and number of anchored and unanchored polymers per cell. This is a pioneering study identifying the role of *drmA* encoding a DUF998 in the acquisition of high-level DAP-R, biofilm formation and the modulation of Lys-Glc_2_-DAG levels in enterococci. Our work indicates a role for DUF998-containing proteins as a new class of lipid regulators with widespread distribution in *Bacteria* and *Archaea*.

## Acknowledgements

This work was supported by R01AI178692 to K.P., F.M., K.D. and Z.G, by R01AI148366 to K.P and Z.G., and by the Cecil H. and Ida Green Chair in Systems Biology Science to K.P. T.N. was supported by the State Department Grant number SGG80022GR0042 provided by the U.S. Embassy in Tbilisi, Georgia. We thank Dr. Tahira Amdid Ratna for training on the biofilm assay and confocal microscopy. We thank Dr. Roberto Jhonatan Olea Ozuna for discussion on statistical analysis of TLC quantification.

---

**Figure S1.**
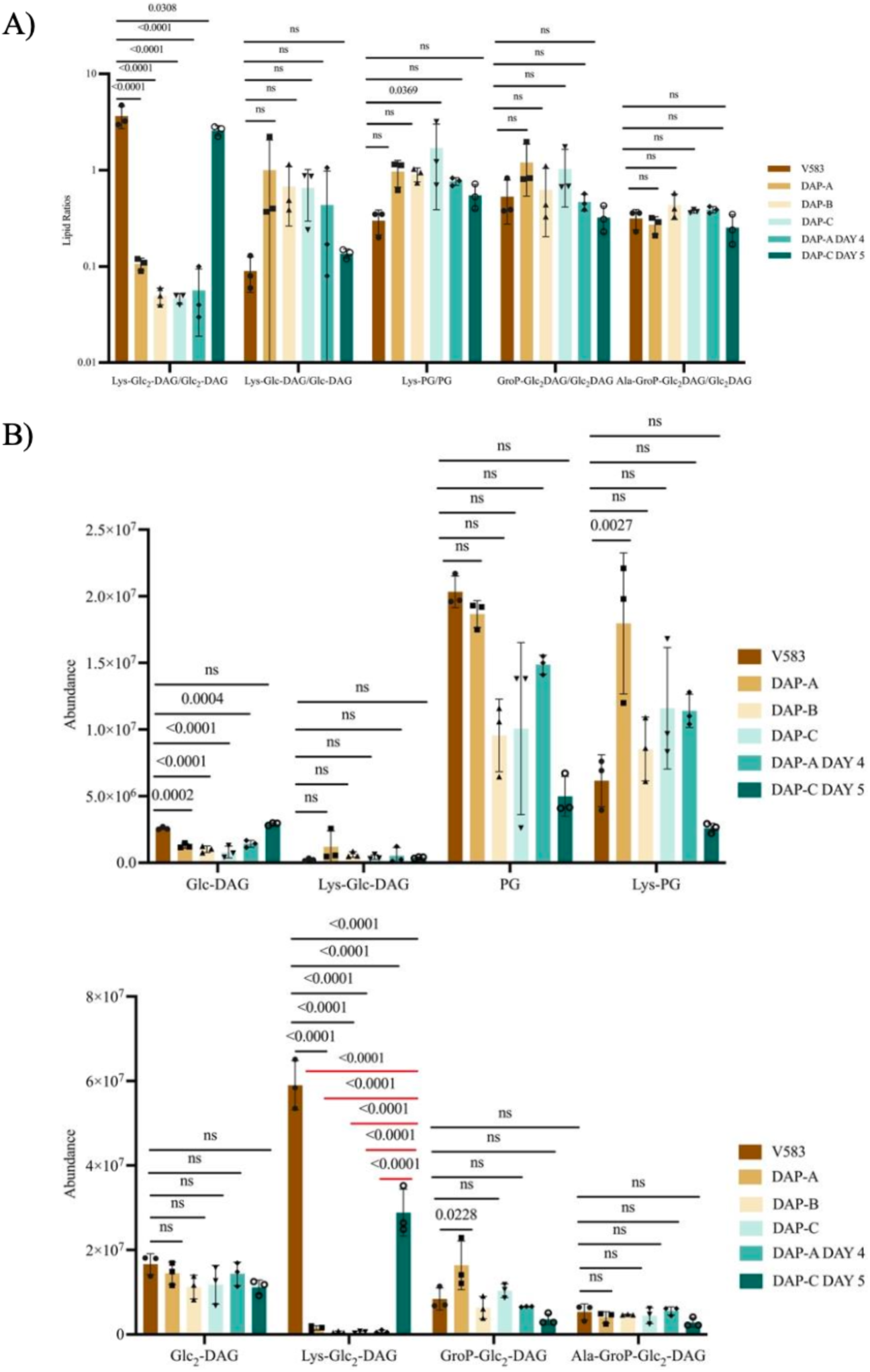
Normalized ratios of lipid substrates and lipid products and absolute intensity of individual lipid indicate significant Lys-Glc_2_-DAG reduction in the DAP-R strains with *drmA* mutations compared to those without. Intensity values for each lysine-modified lipid were normalized to their specific biosynthetic precursor lipid (Figure S1A) and their absolute intensity values with no normalization (Figure S1B) for n=3 independent replicates. Bars on graph represent the average lipid ratio across replicates. Bars represent mean ± standard deviation. Significance was assessed using ordinary one-way ANOVA for Dunnett’s multiple comparisons test. Individual p-values of normalized lipid substrate-to-product ratios are shown in the graphs.

**Figure S2.**
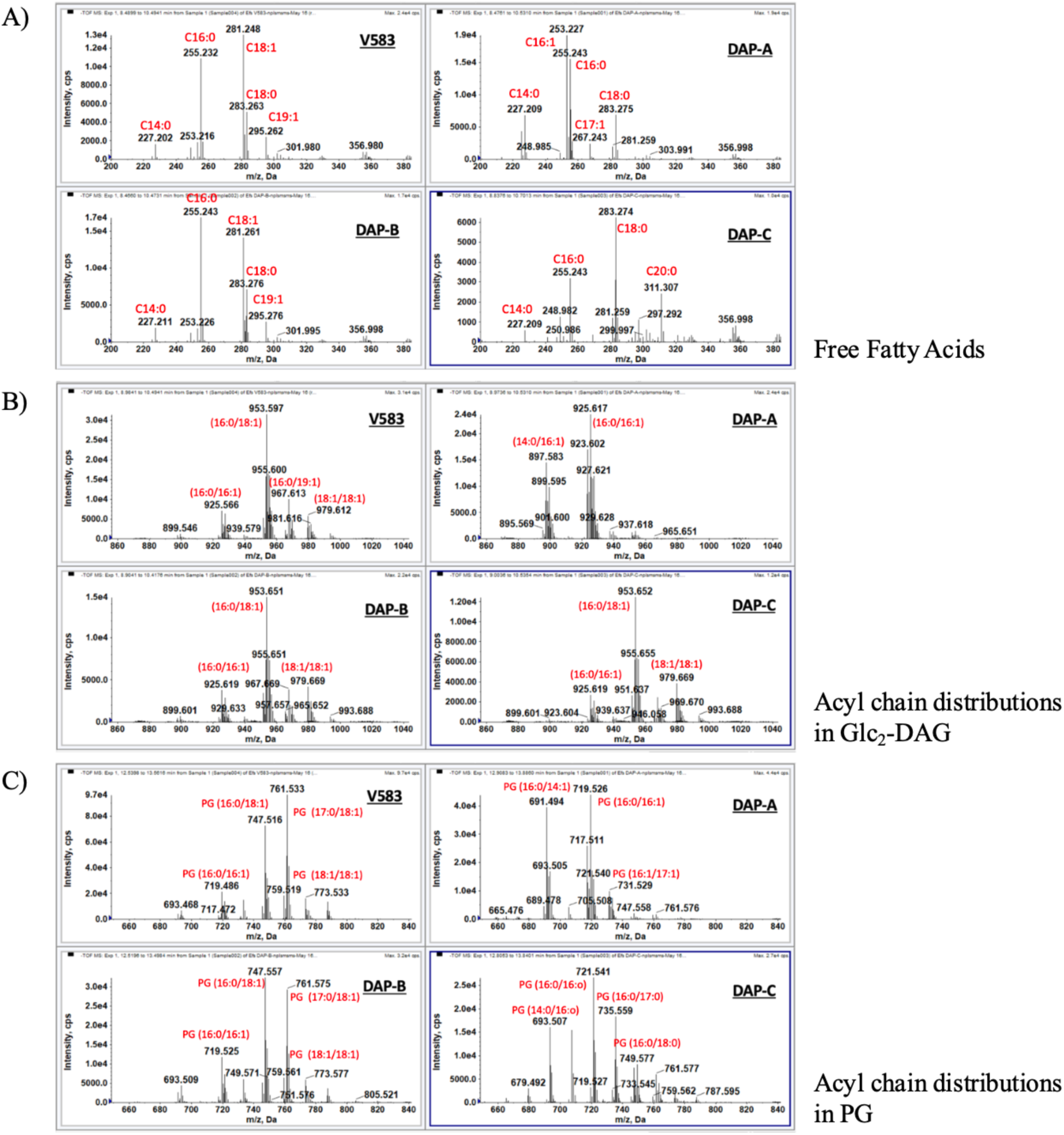
DAP-R strains have differences in fatty acyl composition. The positive ion mass spectrums indicate the parental V583, and the DAP-R strains have alterations in the free fatty acids (A), and in acyl chain distributions of Glc_2_-DAG and PG (B and C). PG – Phosphatidylglycerol; Glc_2_-DAG – Diglucosyl-diacylglycerol.

**Figure S3.**
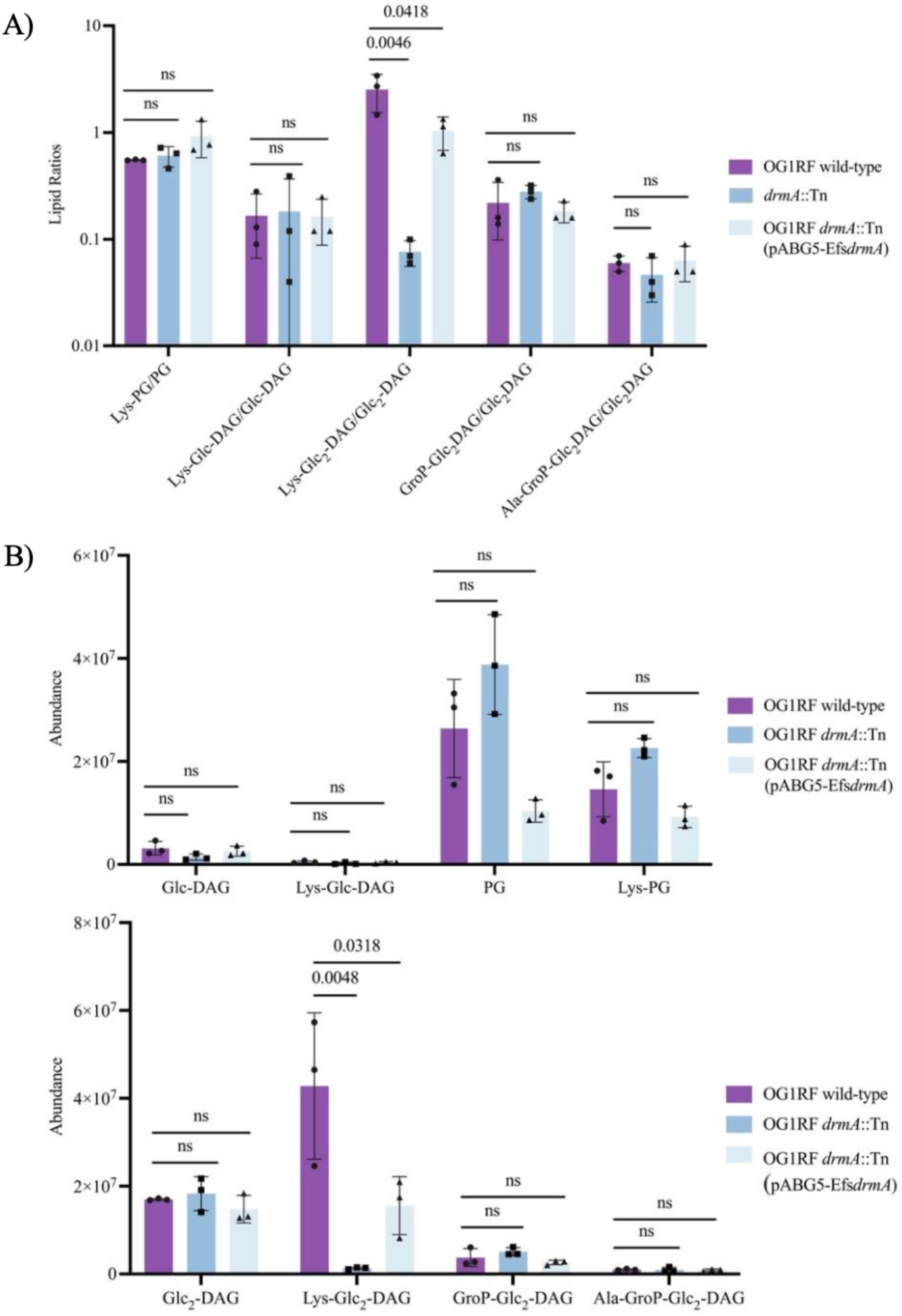
Normalized ratios of lipid substrates and lipid products and absolute intensity of individual lipid indicates significant Lys-Glc_2_-DAG reduction in the *drmA*::Tn compared to *E. faecalis* OG1RF. Intensity values for each lysine-modified lipid were normalized to their specific biosynthetic precursor lipids (Figure S3A) and their absolute intensity values with no normalization (Figure S3B) for n=3 independent replicates. Bars on graph represent the average lipid ratio across replicates. Bars represent the mean ± standard deviation. Significance was assessed using ordinary one-way ANOVA for Dunnett’s multiple comparisons test. Individual p-values of normalized ratios of lipid substrate-to-product ratios are shown in the graphs.

**Figure S4.**
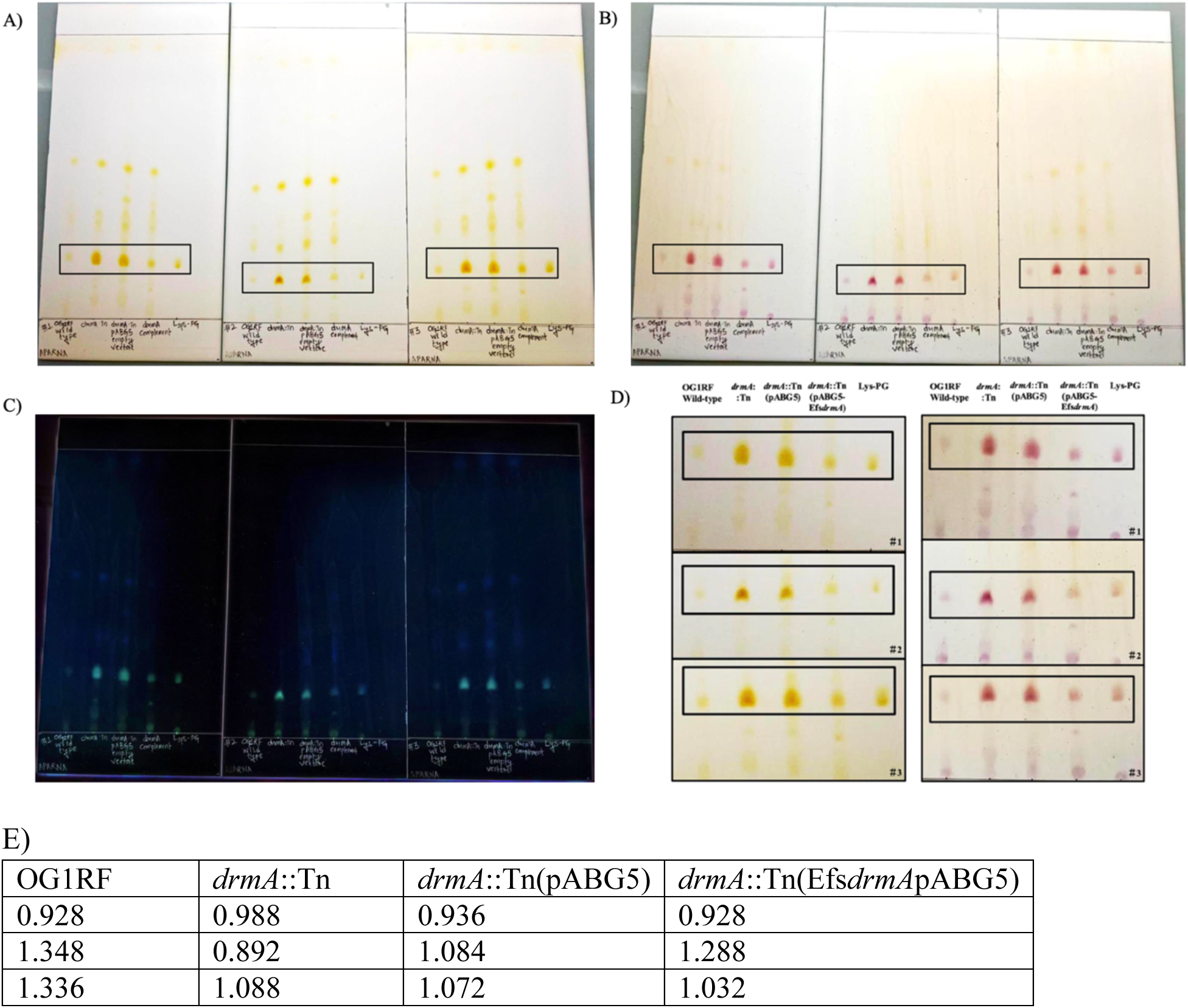
Raw TLC images indicate *E. faecalis* OG1RF *drmA* mutant synthesizes increased Lys-PG levels. The *E. faecalis* strains OG1RF wild-type, *drmA*::Tn, *drmA*::Tn (pABG5) and *drmA*::Tn (pABG5-Efs*drmA*) were spotted on TLC plates in triplicate. The plates were run in a chloroform:methanol:water (65:25:4) solvent system. A) The yellow spots were developed in presence of iodine. B) The TLC plates were sprayed with ninhydrin to detect aminolipids. Avanti Lys-PG 18:1 was used as a lipid standard in all TLC plates. C) The TLC image was inverted in Fiji (Image J) and used for TLC quantification. D) Lys-PG spots across three replicates in iodine and ninhydrin conditions for *E. faecalis* strains OG1RF wild-type, *drmA*::Tn, *drmA*::Tn (pABG5) and *drmA*::Tn (pABG5-Efs*drmA*). E) O.D.600nm values of triplicates per each strain. Cell pellets were obtained from 100 mLs of overnight cultures. These pellets were used for lipid extraction by acidic Bligh-Dyer as mentioned in materials and methods section. 100 μLs of chloroform was used to resuspend the dried lipids. 10 μLs of each lipid extract was used for each spot in TLC.

**Figure S5.**
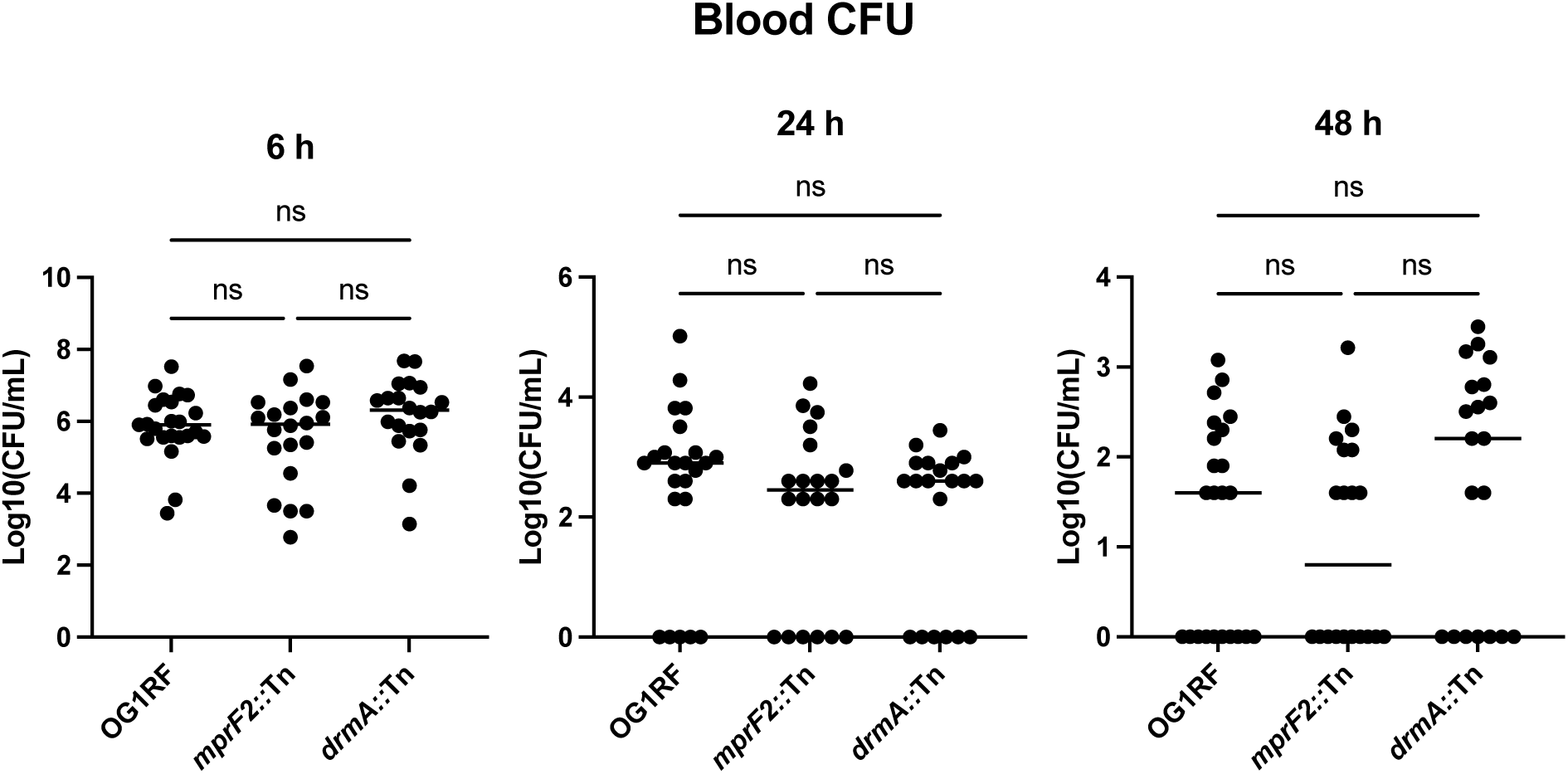
Blood survival of *E. faecalis*. Groups of ∼7 week old male CD-1 mice were injected via tail vein with ∼3.5 - 5 x 10^8^ CFU of OG1RF, Δ*mprf2*, and Δ*drmA* strains and blood was collected at 6, 24 and 48 hour post infection. A decrease in CFU was observed at 48 hpi for mice infected with Δ*mprF2* compared to OG1RF and Δ*drmA*. OG1RF *n =* 23, Δ*mprF2 n =* 20, and Δ*drmA n =* 20. Median indicated. Statistical analyses: Kruskal-Wallis with uncorrected Dunn’s test. ns, not significant.

**Figure S6.**
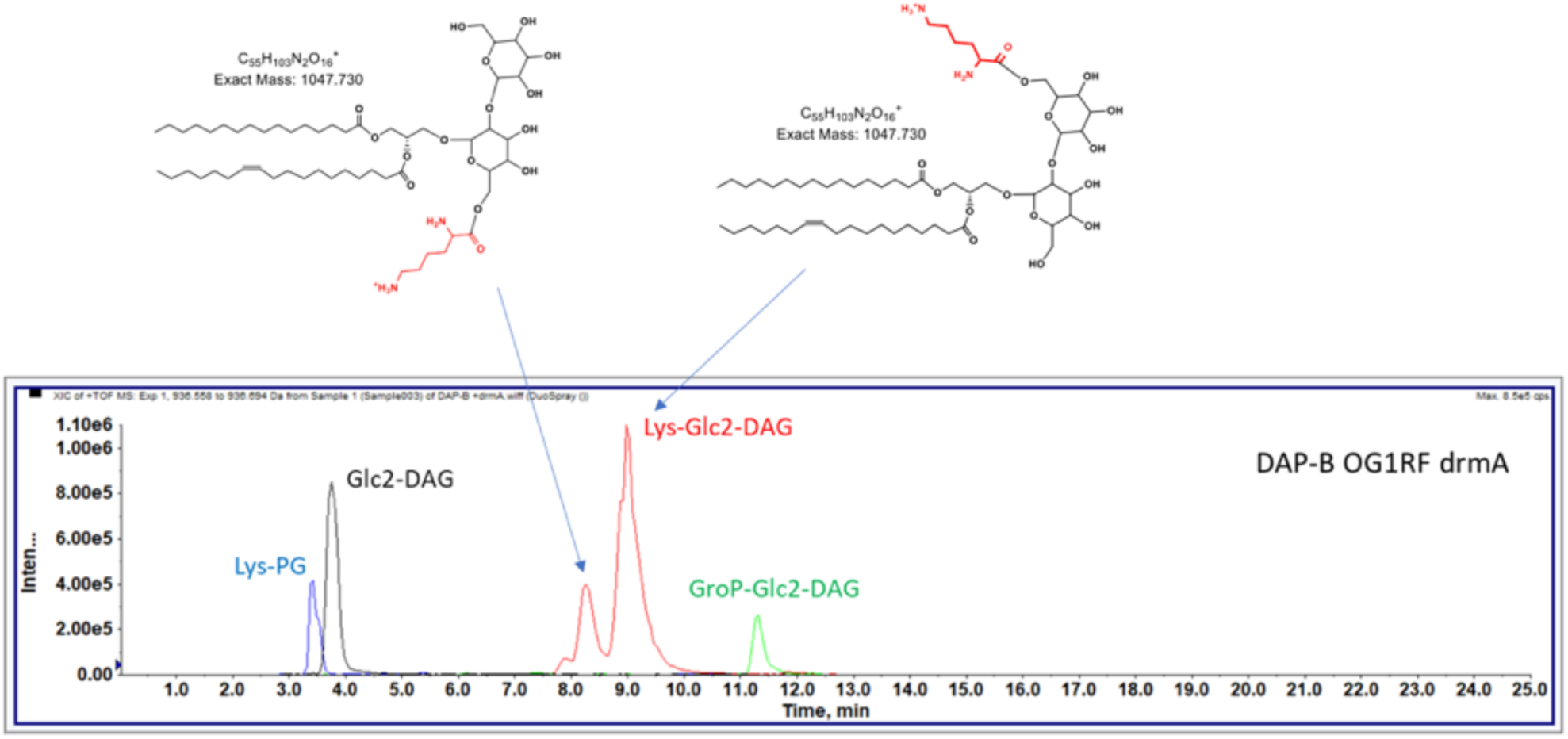
Two distinct lysine linkage positions on Glc_2_-DAG. The ion chromatogram indicates the two Lys-Glc_2_-DAG species with an identical molecular mass (*m/z* 1047.730) but different retention times. The structures shown indicate that lysine is attached to either the proximal or distal glucose residue of Glc₂-DAG. Lys-PG – Lysyl-phosphatidylglycerol; Glc_2_-DAG – Diglucosyl-diacylglycerol; Lys-Glc_2_-DAG – Lysyl-diglucosyl-diacylglycerol; GroP-Glc_2_-DAG – Glycerophospho-diglucosyl-diacylglycerol.

**Figure S7.**
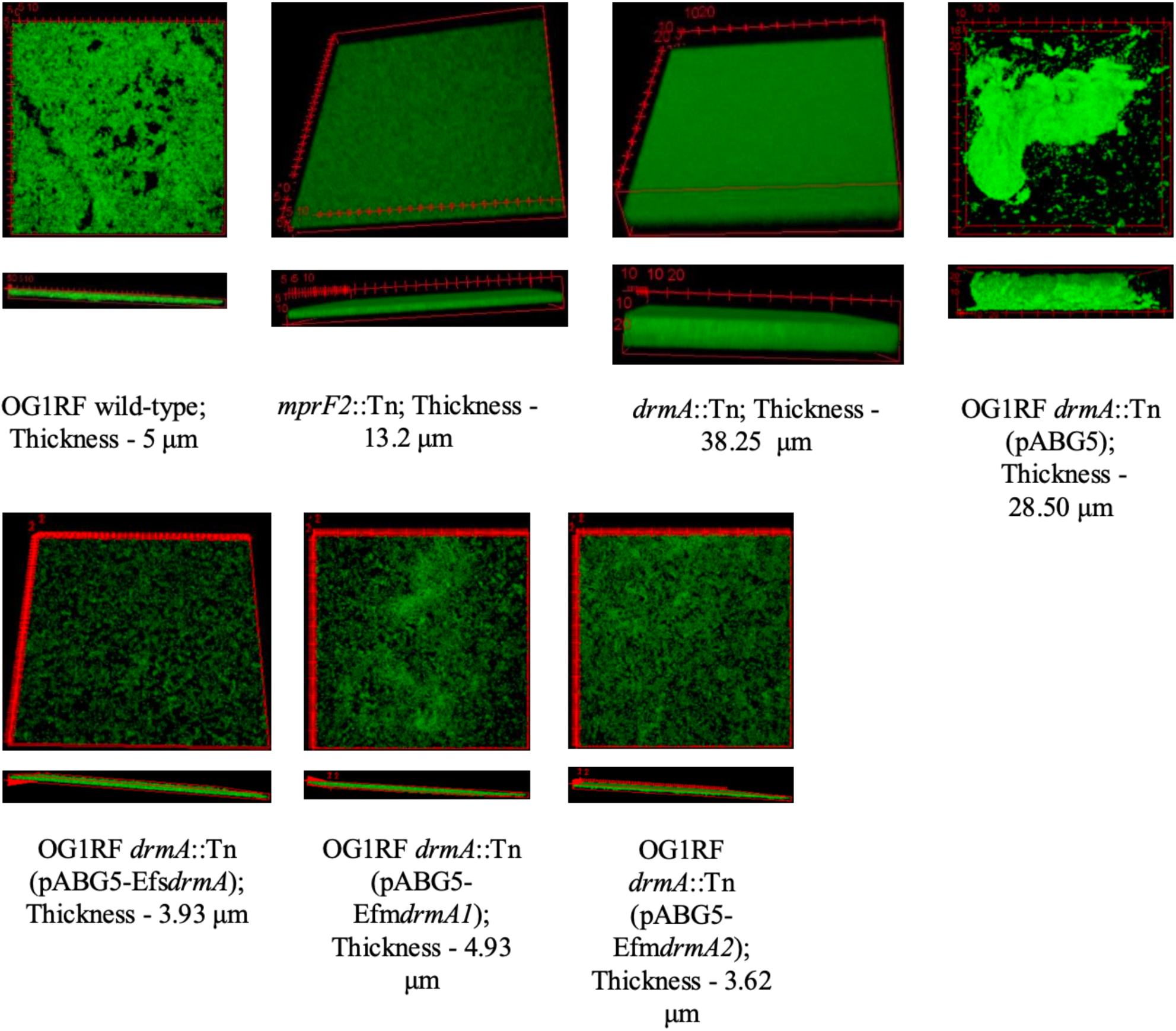
OG1RF *drmA*::Tn forms significantly thicker biofilms. Z-stack images acquired by confocal microscope (Zeiss LSM880) are shown in top-view and side-view orientations. The side-view projections highlight differences in biofilm thickness among the strains.

**Figure S8.**
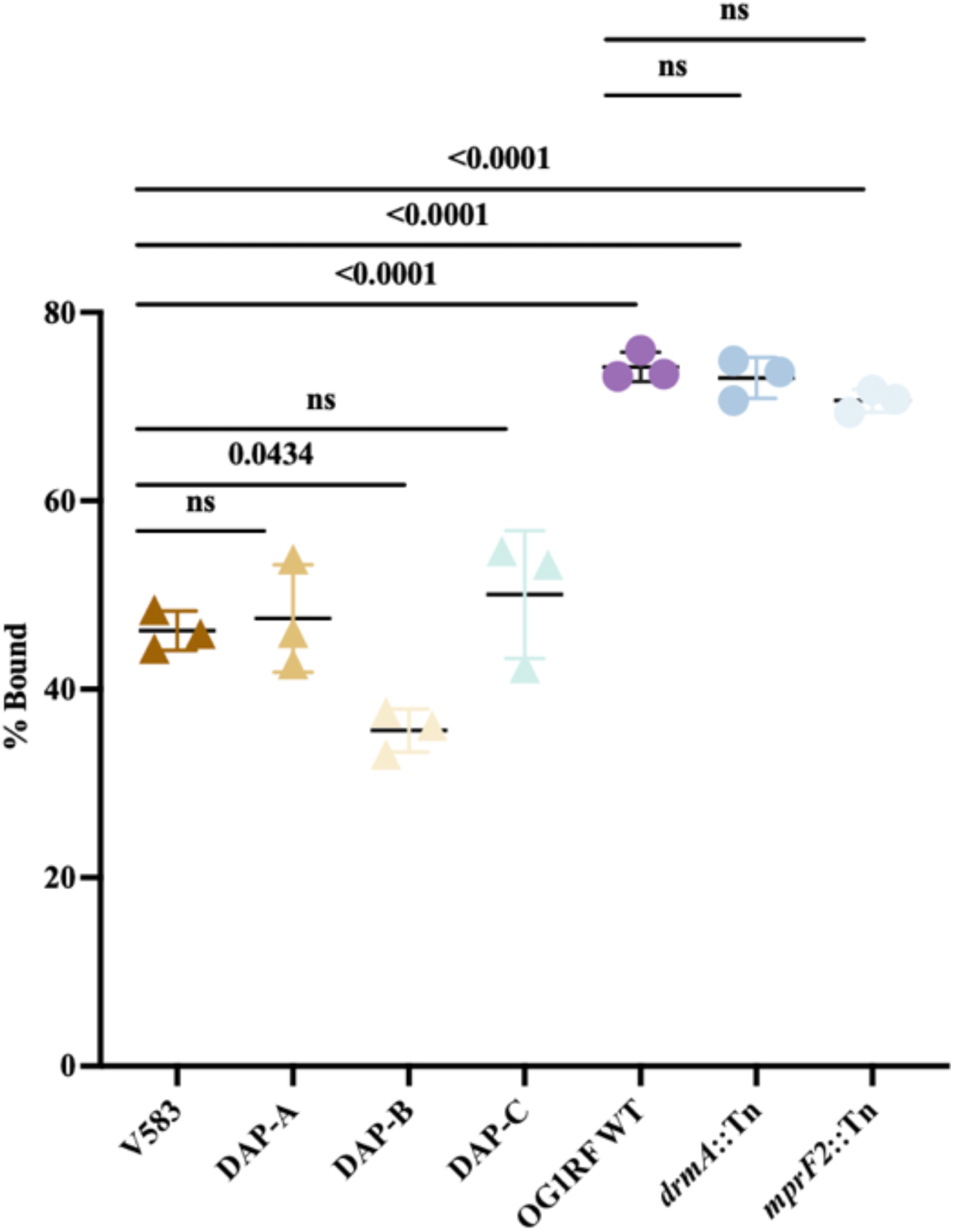
Enterococcal strains may possess compensatory mechanism(s) to regulate membrane charge. The y-axis shows the percentage of cytochrome *c* bound (% bound), and the x-axis shows the different strains: V583, DAP-A, DAP-B, DAP-C, OG1RF wild type, *drmA::Tn*, and *mprF2::Tn*. Data points represent individual measurements, and symbols indicate the mean ± standard deviation for each strain. Significance was assessed using ordinary one-way ANOVA with Tukey’s multiple comparisons test.

**Table S1:** *drmA* mutations identified in previous studies of daptomycin resistance in *Enterococcus faecalis*.

| Parental Strain | *drmA* gene locus ID | Derivative strain | Substitution/Mutation | DAP MIC (µg/ml) | Source |
| --- | --- | --- | --- | --- | --- |
| *E. faecalis* V583 | EF1797 | DAP-A | D30fs | 256 | (1) |
| *E. faecalis* V583 | EF1797 | DAP-B | V16insIS256 | 256 | (1) |
| *E. faecalis* V583 | EF1797 | DAP-C | G130_F154del | 512 | (1) |
| *E. faecalis* V583 | EF1797 | DAP-A Day 4 | D30fs | 16 | (1) |
| *E. faecalis* S613 | - | TDR13 | L4stop | - | (2) |
| *E. faecalis* S613 | - | TDR22 | N150fs | - | (2) |
| *E. faecalis* S613 | - | TDR28 | N150fs | - | (2) |
| *E. faecalis* OG1RF | OG1RF_11507 | DAP21 | Frameshift 19 | 128 | (3) |
| *E. faecalis* OG1RF | OG1RF_11507 | DAP22 | Nonsense L4 | 128 | (3) |
| *E. faecalis* AR01/DGVS | CG806_RS08810 | 2AD | L4*STOP truncation | 128-256 | (5) |
| *E. faecalis* AR01/DGVS | CG806_RS08810 | 3AD | G127D substitution | 128-256 | (5) |
| *E. faecalis* AR01/DGVS | CG806_RS08810 | 5AD | Premature stop codon downstream at residue 183 | 128 | (5) |
| *E. faecalis* JH2-2 | O994_RS07740 | 3JD | Frameshift 36 | 128 | (5) |
| *E. faecalis* JH2-2 | O994_RS07740 | 4JD | L4*STOP truncation | 128 | (5) |
| *E. faecalis* JH2-2 | O994_RS07740 | 5JD | G62R substitution | 128 | (5) |
| *E. faecalis* (ATCC 700802) | EF1797 | Efs807 pass (Day 15 surotomycin serial passage isolate) | N150fs | 32 | (6) |

